# The binding sites of E2F transcription factor in *Drosophila* metabolic genes are functionally distinct

**DOI:** 10.1101/2022.11.22.517506

**Authors:** Maria Paula Zappia, Yong-Jae Kwon, Anton Westacott, Isabel Liseth, Hyun Min Lee, Abul B.M.M.K. Islam, Jiyeon Kim, Maxim V. Frolov

## Abstract

The canonical role of the transcription factor E2F is to control the expression of cell cycle genes by binding to the E2F sites in their promoters. However, the list of putative E2F target genes is extensive and includes many metabolic genes, yet the significance of E2F in controlling expression of these genes remains largely unknown. Here, we used the CRISPR/Cas9 technology to introduce point mutations in the E2F sites upstream of five endogenous metabolic genes in *Drosophila*. We found that the impact of these mutations on both the recruitment of E2F and the expression of the target genes varied, with the glycolytic gene, *Phosphoglycerate kinase* (*Pgk)*, being mostly affected. The loss of E2F regulation on *Pgk* gene led to a decrease in glycolytic flux, TCA cycle intermediates levels, ATP content and an abnormal mitochondrial morphology. Remarkably, chromatin accessibility was significantly reduced at multiple genomic regions in *Pgk*^Δ*E2F*^ mutants. These regions contained hundreds of genes, including metabolic genes that were downregulated in *Pgk*^Δ*E2F*^ mutants. Moreover, *Pgk*^Δ*E2F*^ animals had shortened life span and exhibited defects in high-energy consuming organs, such as ovaries and muscles. Collectively, our results illustrate how the pleiotropic effects on metabolism, gene expression and development in the *Pgk*^Δ*E2F*^ animals underscore the importance of E2F regulation on a single E2F target, *Pgk*.

## INTRODUCTION

The E2F transcription factor is a heterodimeric complex between one E2F and one DP subunits, which has been initially characterized several decades ago by its binding to a short 8 bp degenerative sequence in the adenovirus E2 promoter. These, so called E2F-binding sites, are also present in the promoters of many cell cycle genes and mediate their transcriptional activation in a cell cycle dependent manner. The activity of the E2F transcription factor is negatively regulated by the Retinoblastoma tumor suppressor protein (pRB) that binds to E2F and masks its transactivation domain. The inhibition of E2F by pRB is released in late G1, as cyclin dependent kinases phosphorylate pRB, thus dissociating E2F-pRB complexes and releasing free E2F that activates the transcriptional program for G1-S transition and drives the entry to S phase (Dyson, 1998, 2016; Rubin *et al*, 2020; Weinberg, 1995; Blais & Dynlacht, 2004). This elegant, yet simple, view of E2F and pRB provides a satisfactory explanation of why the functional inactivation of pRB makes cells insensitive to growth inhibitory signals and, therefore, helps to rationalize frequent mutations of the Rb pathway in cancer.

The model of how pRB regulates E2F-dependent transcription served as a framework for follow up studies that revealed a much more complex picture. Unexpectedly, the list of E2F target genes extends far beyond cell cycle genes. Numerous analyses on genome-wide expression and occupancy profiles in various systems revealed thousands of E2F-regulated genes that are involved in almost every cellular function (Muller *et al*, 2001; Julian *et al*, 2015; Denechaud *et al*, 2016; Georlette *et al*, 2007; Korenjak *et al*, 2012; Dimova *et al*, 2003; Zappia *et al*, 2019). Among E2F targets are genes involved in differentiation programs, metabolism, mitochondrial function, apoptosis and others. Many of these transcriptional programs appear to be cell type specific illustrating that E2F may have unique functions in different contexts (Chicas *et al*, 2010; Rabinovich *et al*, 2008). Collectively, these observations support the idea that E2F-dependent transcription is a key aspect of E2F function and, thus, highlight the significance of *cis* regulating elements, including E2F sites, that mediate the recruitment of E2F to its target genes.

Given that E2F regulates thousands of genes, one of the important questions in the field is how many among them are the key effector genes. Is the function of E2F the net result of E2F controlling hundreds of genes with either many of them being important or only a few being rate limiting? Earlier studies in both flies and mammals showed that the overexpression of *cyclin E* can rescue defects in S phase entry in E2F-deficient cells (Duronio *et al*, 1995; Lukas *et al*, 1997), thus, leading to the idea that *cyclin E* is the key downstream target of E2F. The other key target is thought to be *string*, encoding Cdc25, which can drive E2F mutant cells through G2/M transition (Neufeld *et al*, 1998). One limitation of these experiments is that genes are overexpressed at non-physiologically high levels and that the rescue is incomplete as cells are unable to sustain normal proliferation. Later genetic experiments demonstrated that the list of key cell cycle targets is likely to be much larger, as haploinsufficiency for half of E2F-target genes modified the E2F-dependent phenotype (Herr *et al*, 2012). Although these experiments provided a glimpse at the potential rate-limiting E2F targets, they did not address the functional importance of E2F binding sites in their promoters. Another complexity arises from E2F participating in multiple tissue-specific transcriptional programs. Thus, the relative importance of E2F targets likely varies between different cell types and, therefore, this question cannot be easily addressed in cell lines.

Model systems, such as *Drosophila*, which has a streamlined and highly conserved version of the Rb pathway, are advantageous in exploring context dependency of E2F- dependent transcription during animal development (van den Heuvel & Dyson, 2008). In flies, there are two E2F genes, *E2f1* and *E2f2*, a single *Dp* gene and an Rb ortholog, *Rbf*. Since Dp is required for both E2Fs to bind to DNA, inactivation of Dp either by RNAi or by genetic mutations has been often used to explore the consequence of the loss of E2F regulation (Frolov *et al*, 2005; Royzman *et al*, 1997). Surprisingly, *Dp* deficient animals develop relatively normally until late pupal stages when the lethality occurs due to defects in metabolic tissues, such as muscle and fat body (Guarner *et al*, 2017; Zappia & Frolov, 2016a). A common feature in both Dp-deficient tissues is a change in carbohydrate metabolism, which, at least partially, accounts for animal lethality (Zappia *et al*, 2021). Integrative analysis of proteomic, transcriptomic, and ChIP-seq data suggest that E2F/Dp/Rbf directly activates the expression of glycolytic and mitochondrial genes during muscle development (Zappia *et al*, 2019). Given the importance of this transcriptional program, and that many of these metabolic genes contain E2F binding sites, we aimed to study how these genes are regulated by E2F.

Using CRISPR/Cas9 technology in *Drosophila*, we systematically edited putative E2F binding sites in five metabolic genes. We determined the impact of these mutations on the recruitment of E2F/Dp/Rbf to the regulatory regions of these genes, the expression of the target genes and the impact on fly physiology. We found a wide range of effects, which were gene specific. For most target genes, E2F appears to provide a limited contribution to their expression. In contrast, E2F regulation was particularly important for the glycolytic gene *Phosphoglycerate kinase* (*Pgk*), as the mutations on E2F sites resulted in a strong reduction in *Pgk* expression, mimicked several aspects of the *Pgk* mutant phenotype, and exerted broad effect on development. Strikingly, the animals showed low levels of glycolytic and tricarboxylic acid (TCA) cycle intermediates, defects in mitochondria structure and compromised muscle function. Unexpectedly, a widespread reduction in chromatin accessibility was detected and it correlated with a broad decrease in gene expression, thus supporting the notion that changes in the levels of some metabolites can lead to changes in the epigenetic landscape. Collectively, our data illustrate how the loss of E2F regulation on a single metabolic gene, such as *Pgk*, can indirectly impact the expression of other genes, including E2F targets, and suggest that E2F dependent regulation is a complex combination of both direct and indirect effects.

## RESULTS

### E2F binding sites contribute to the transcriptional activation of metabolic genes in reporter assays *in vitro*

E2F/Dp has an essential role in adult skeletal muscle that explains the lethality of E2F- deficient animals (Zappia & Frolov, 2016a). Unlike its canonical role in the regulation of the cell cycle genes, E2F/Dp together with Rbf directly activate the expression of metabolic genes during late stages of muscle development (Zappia *et al*, 2019). To understand the mechanisms by which E2F regulates this process, we examined the function of the *cis* regulating elements located on the vicinity of E2F-target genes that mediate the recruitment of E2F/Dp/Rbf. We took advantage of available proteomic and transcriptomic profiles for Dp- and Rbf-deficient muscles (Zappia *et al*, 2021), respectively, as well as muscle-specific ChIP-seq for both Dp and Rbf (Zappia *et al*, 2019). Since the loss of Dp results in severe changes in carbohydrate metabolism we first focused on analyzing genes encoding for proteins involved in glucose metabolism (Figure 1A), in which the protein levels were reduced in Dp-depleted muscles (Figure 1B, (Zappia *et al*, 2021)). Then, from this list we selected genes that are downregulated upon Rbf-depletion using the RNA-seq dataset for Rbf-deficient muscles and define a set of genes in which both Dp and Rbf are needed for their transcriptional activation (Figure 1C, (Zappia *et al*, 2019)). Finally, we mined ChIP-seq for Dp and Rbf from skeletal muscle to identify genes with closely matching summits of both Dp and Rbf peaks in the vicinity of their transcription start sites (TSS, (Zappia *et al*, 2019)). Using these three stringent criteria, we selected three glycolytic genes, *Aldolase* 1 (*Ald1*), *Phosphoglycerate kinase* (*Pgk*) and *Pyruvate kinase* (*Pyk*), and one gene linking glycolysis to triglyceride synthesis, *Glycerol-3-phosphate dehydrogenase 1* (*Gpdh*) (Figure 1A). To broaden the list of candidates, we also included the citrate synthase, *knockdown* (*kdn*), of the tricarboxylic acid (TCA) cycle and *Cytochrome c proximal* (*Cyt-c-p*) of the electron transport chain in our analysis (Figure 1A). In sum, all selected genes encoding for metabolic enzymes are strongly downregulated in both Dp- and Rbf- depleted muscles and have closely matching Dp and Rbf peaks in the vicinity of their TSSs (Figure 1E-J), thus implying that E2F could be directly activating the expression of these genes.

**Figure 1:**
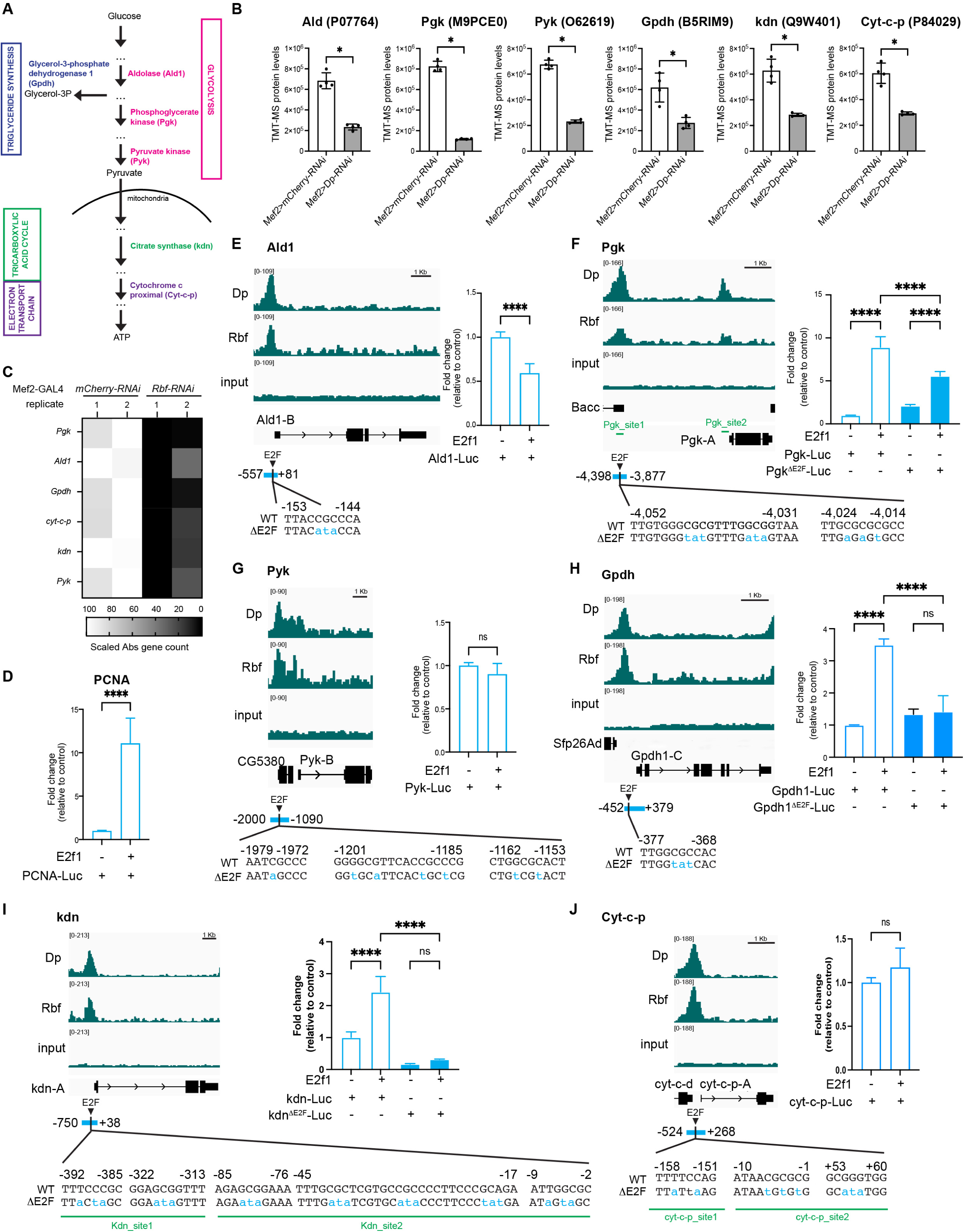
E2F target genes involved in metabolism. (A) Simplified illustration of the enzymes Aldolase (Ald1), Phosphoglycerate kinase (Pgk), Pyruvate kinase (Pyk) in the glycolytic pathway, Glycerol-3-phosphate dehydrogenase 1 (Gpdh) connecting triglyceride synthesis with glycolysis, knockdown (kdn) in the TCA cycle, and cytochrome c proximal (Cyt-c-p) in the electron transport chain. (B) Protein levels quantified by TMT-MS in flight muscles of *Mef2>mCherry-RNAi* and *Mef2>Dp-RNAi*, scatter plot with bar showing Mean ± SD, Mann-Whitney test, * p<0.05, n=4 (C) Transcripts levels measured by RNAseq in *Mef2>mCherry-RNAi* and *Mef2>Rbf-RNAi*, heatmap depicting expression of selected genes only. Abs count was normalized and scaled, n=2 samples per genotype (D-J) Left panel: ChIPseq for Rbf, Dp and the control input visualized with Integrative Genomics Viewer browser for the genomic regions surrounding the genes. The most predominantly expressed transcript in adult flies, based on the FlyAtlas 2 profile (Leader *et al*, 2018), is displayed; n = 2 samples/condition. Read scales and genomic scales included on top left and top right, respectively. GroupAuto scale was used. Right panel: Dual luciferase reporter assay performed in S2R+ cells for (D) *PCNA-Luc*, (E) *Ald1-Luc*, (F) *Pgk-Luc*, (G) *Pyk-Luc*, (H) *Gpdh-Luc*, (I) *kdn-Luc*, and (J) *Cyt-c-p-Luc*. Values for Firefly Luciferase luminescence were normalized to Renilla luminescence. Ratio was plotte as fold change relative to control (no E2f1 expression). Mutated E2F sites are indicated as ΔE2F. Bar plot showing Mean ± SD, (F, H, I) One-way ANOVA followed by Tukey’s multiple comparisons test, (D, E, G, J) Unpaired t- test, ns p>0.05, **** p< 0.0001, n=2 replicates per group, N=2 independent experiment. Bottom panel: E2F binding sites identified using degenerated motif WKNSCGCSMM. Mutations on the core are indicated in blue as ΔE2F. Blue bars indicate regions amplified and cloned upstream luciferase reporter. Positions are relative to TSS. Green bars indicate different sites amplified by ChIP-qPCR in Figure 2 and Figure S1A. Full genotypes: (B) *w-; Mef2-GAL4/UAS-mCherry-RNAi* (white bar) and *w-; UAS-Dp-RNAi; Mef2-GAL4* (grey bar) (C) *w-,UAS-Dicer2; +; Mef2-GAL4/UAS-mCherry-RNAi*, and *w-,UAS- Dicer2; +; Mef2-GAL4/UAS-Rbf-RNAi*

Next, we analyzed the sequences encompassing the Dp and Rbf summits to identify putative E2F sites. We used the degenerative motif WKNSCGCSMM that was previously identified by *de novo* discovery motif as an E2F site in the flight muscles (Zappia *et al*, 2019). For each gene, a single or multiple predicted E2F sites were found (Figure 1E-J). As an initial test for the functionality of these sites, we generated six luciferase constructs containing the corresponding genomic regions for each gene (visualized as a blue square) and tested the ability of a sole *Drosophila* E2F activator, E2f1, to activate these reporters in S2R+ cells. The luciferase constructs along with Renilla reporter, which was used to normalize transfection efficiency, were transiently transfected in S2R+ cells with or without the E2f1 expression plasmid. Luciferase activity was measured using a dual luciferase reporter assay. As expected, the transfection of the E2f1 expression plasmid led to a 10-fold activation of the *PCNA-luc* reporter, a well-known E2F reporter (Figure 1D) (Sawado *et al*, 1998). In contrast, the generated luciferase constructs responded differently to the same amount of transfected E2f1. Three luciferase constructs, *Pgk-luc*, *Gpdh-luc* and *kdn-luc*, were activated 3 to 10-fold by E2f1 and largely matched the activity of the *PCNA-luc* reporter (Figure 1 D,F,H,I). To determine whether E2f1-dependent activation is mediated through E2F binding sites, point mutations in the core of the E2F motif for each predicted E2F site were introduced in each of the luciferase reporters (marked in blue letters in Figure 1F,H,I). The *GpdhΔE2F-Luc* and *kdnΔE2F-Luc* reporters no longer responded to E2f1 (Figure 1H,I). Mutating E2F binding sites in the *PgkΔE2F-luc* significantly reduced but did not completely abrogate the ability of E2f1 to activate the reporter (Figure 1F). Unlike *Pgk-luc*, *kdn-luc* and *Gpdh-luc*, the luciferase reporters for *Ald1*, *Cyt-c-p* and *Pyk* failed to be activated by E2f1, as the luciferase activities were largely indistinguishable with or without the transfected E2f1 expression plasmid (Figure 1 E, G, J). Overall, these data indicate that E2F regulates the transcription of several metabolic genes, including *Gpdh*, *Pgk* and *kdn*, *in vitro*.

### The E2F binding sites recruit the transcriptional complex E2F/Dp/Rbf for full activation of gene expression *in vivo*

To understand the role of the newly identified E2F binding sites in the regulation of the endogenous metabolic genes *in vivo*, we introduced precise point mutations in the core elements of E2F motifs *via* genome editing using CRISPR/Cas-9 -Catalyzed Homology-Directed Repair. We generated flies carrying such mutant alleles. For each gene, the mutated sequences are shown in blue in Figure 1 E-J. For *Pgk*, *Gpdh* and *kdn*, the point substitutions are identical to the mutated sequences of the luciferase reporters *PgkΔE2F-Luc*, *GpdhΔE2F-Luc* and *kdnΔE2F- Luc* (Figure 1 F,H,I). For the exception of *Pyk*, we successfully generated *Gpdh*^ΔE2F^, *Pgk^ΔE2F^*, *kdn*^ΔE2F^, *Ald1^ΔE2F^* and *Cyt-c-p*^ΔE2F^ mutant alleles. Each of these alleles contained the abovementioned mutations in E2F sites that was confirmed by sequencing. In order to eliminate off-target effects of CRISPR/Cas-9 gene editing, all ΔE2F site-edited lines were backcrossed *en masse* to the wild-type strain *w^1118^* for six consecutive generations. Individual males from the sixth generation were used to establish at least two independent lines for each allele. The *Gpdh*^ΔE2F^, *Pgk*^ΔE2F^, *kdn*^ΔE2F^, *Ald1*^ΔE2F^ and *Cyt-c-p*^ΔE2F^ alleles were homozygous viable.

To determine the effect on the recruitment of E2F/Dp/Rbf in response to mutating E2F sites, we performed ChIP-qPCR using anti-Dp and anti-Rbf antibodies. A known E2F target gene, *Arp53D*, was used as a positive internal control to assess the level of Dp and Rbf enrichment in each chromatin immunoprecipitation (Figure 2A-H, S1A). Chromatin was prepared from third instar larva of the wild type, *w^1118^*, along with *Gpdh*^ΔE2F^, *Pgk*^ΔE2F^, *kdn*^ΔE2F^, *Ald1*^ΔE2F^ and *Cyt-c-p*^ΔE2F^ homozygous mutant animals, and incubated with anti-Dp, anti-Rbf and non-specific antibodies. The occupancy of both Dp and Rbf at each gene was determined by qPCR using primers flanking the mutated sequences. Remarkably, with the exception of *Cyt-c-p*^ΔE2F^, the recruitments of both Dp and Rbf were severely reduced when E2F sites were mutated (Figure 2A-H, S1A). We noticed that the extent of reduction in the occupancies of Dp and Rbf varied among the mutant alleles. For example, there was a 3-to-4-fold reduction in Dp and Rbf recruitment to the *Gpdh*^ΔE2F^ and *kdn*^ΔE2F^ alleles while mutating E2F sites in *Ald* and *Pgk* resulted in much stronger effects. Remarkably, the binding of Dp and Rbf were completely lost in *Pgk*^ΔE2F^ as the enrichments for Dp and Rbf at *Pgk_Site1* were indistinguishable from that of negative control site (NS) (Figure 2C-D). Notably, the binding to *Pgk_Site2*, another E2F target site in the vicinity of *Pgk-A* TSS that is about 4 kb downstream from where mutations were introduced (Figure 1F), and the binding to *Ald1*, an E2F target site for another glycolytic gene, were not affected (Figure 2C-D). Thus, our data suggest that the engineered mutations prevent the recruitment of E2F/Dp/Rbf in a highly specific manner.

**Figure 2:**
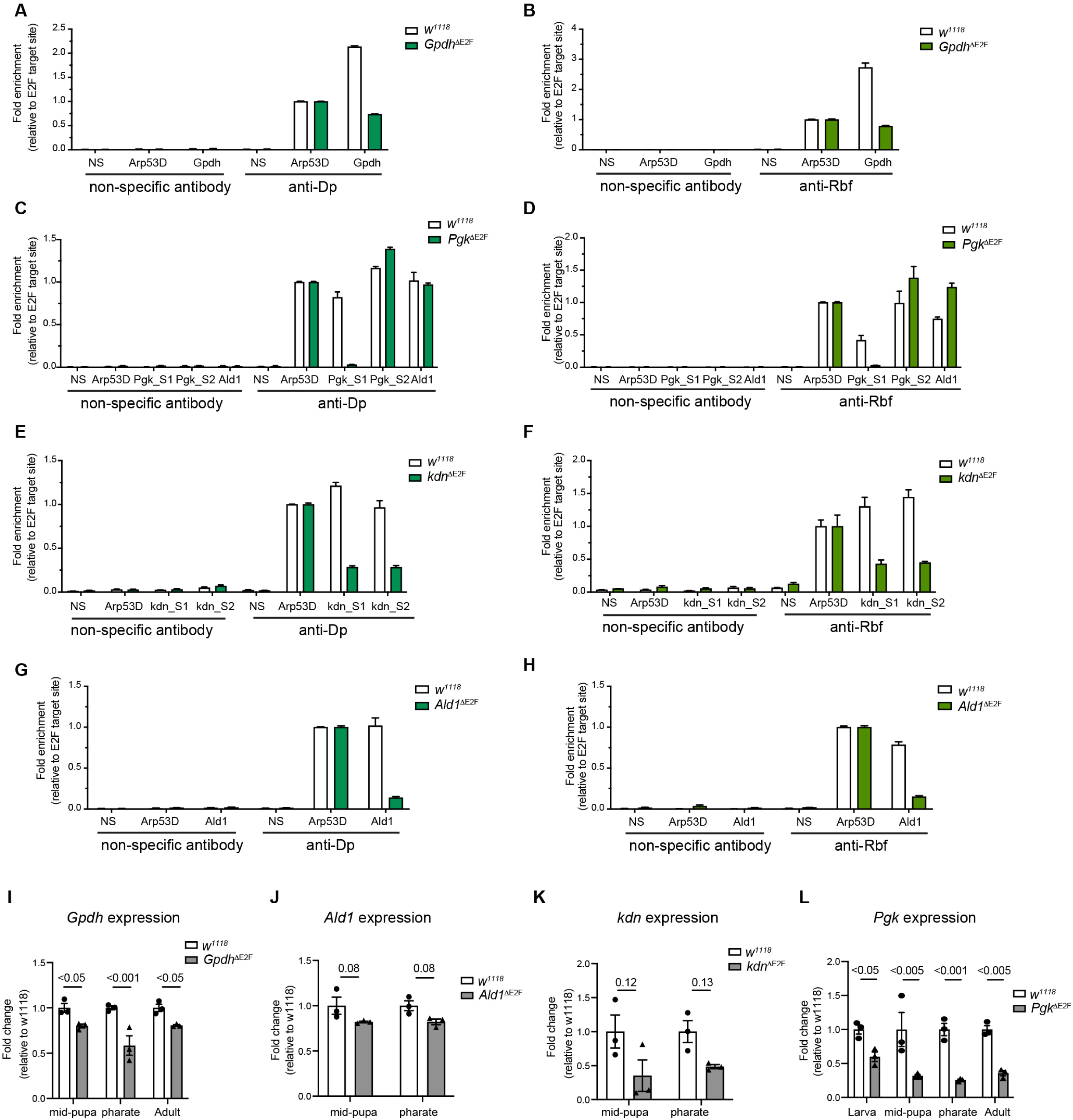
The requirement of the E2F binding sites for the recruitment of E2F/Dp/Rbf and gene expression regulation *in vivo*. (A-H) Dp and Rbf proteins are enriched upstream several metabolic genes in a E2F-dependent manner. Chromatin from third instar larvae was immunoprecipitated with antibodies against Dp (left panel) and Rbf (right panel) and compared with nonspecific control antibodies (IgG on the left panel and anti-Myc on the right). Recruitment was measured flanking the E2F binding sites of the following metabolic genes (A-B) *Gpdh*, (C-D) *Pgk*, (E-F) *kdn*, and (G-H) *Ald1*. The negative site (NS) does not contain predicted E2F-binding sites. The qPCR data are shown as bar plot showing Mean ± SD fold enrichment relative to the positive site, *Apr53D*, for each ChIP sample. Mean ± SEM, N=2 independent experiment. Genomic location of primers amplifying sites S1 and S2 are indicated in Figure 1. (I-L) The expression of the metabolic genes (I) *Gpdh*, (J) *Ald1*, (K) *kdn*, and (L) *Pgk* was measured by RT-qPCR in whole animals staged at third instar larva, mid pupa (48h APF), pharate (96h APF) and 1 -day-old adults. Gene expression was normalized to RpL32 and RpL30 and displayed as fold change relative to control. Scatter dot plots with bars show Mean ± SEM, N= 3 independent samples per group, multiple unpaired t-tests followed by corrected FRD method (Benjamini and Yekutieli), q-value for each comparison is indicated in the plot. Full genotypes: (A-L) control *w^1118^*, (A-B, I) *w^1118^*; *Gpdh*^ΔE2F^;+, (C-D, L) *w^1118^*;*Pgk*^ΔE2F^ line 12;+, (E-F, K) *w^1118^*, *kdn*^ΔE2F^;+ ;+, (G-H, J) *w^1118^*;+;*Ald1*^ΔE2F^

To determine how the changes in the recruitment of E2F/Dp/Rbf impacts the expression of the endogenous genes, RNA was isolated from *w^1118^* control and from homozygous mutant *Gpdh*^ΔE2F^, *Pgk*^ΔE2F^, *kdn*^ΔE2F^, *Ald1*^ΔE2F^ and *Cyt-c-p*^ΔE2F^ animals that were staged at mid-pupa (44h after pupa formation) and pharate (96h after pupa formation). The expression of the corresponding genes was determined by RT-qPCR. There was no reduction in *Cyt-c-p* expression in *cyt-c-p*^ΔE2F^ (Figure S1B), which was expected given that recruitment of Dp was not affected (Figure S1A). In the *Gpdh*^ΔE2F^ and *Ald1*^ΔE2F^ lines, the expression of *Gpdh* and *Ald1* genes were slightly reduced respectively, even though the changes were not statistically significant for Ald1 (Figure 2I-J). The level of the *kdn* mRNA showed a greater reduction in its expression in the *kdn*^ΔE2F^ line. However, it was accompanied by larger variation among the replicates and, therefore, the difference in the expression of *kdn* between the wild type and the *kdn*^ΔE2F^ line was not statistically significantly (Figure 2K).

The strongest effect on gene expression was observed in the *Pgk*^ΔE2F^ line. As shown in Figure 2L, the expression of *Pgk* was reduced several folds in *Pgk*^ΔE2F^ at both mid-pupal and pharate stages. We expanded the analysis and examined the *Pgk* expression at larval and adult stages and found it to be similarly reduced (Figure 2L), thus suggesting that mutating E2F binding sites affects the level of *Pgk* mRNA throughout development. We confirmed that *Pgk* expression was reduced using another, independently generated, *Pgk*^ΔE2F^ line (line 23, Figure S1C) (for details see: Materials and Methods). To exclude the possibility that mutations on the E2F site in *Pgk*^ΔE2F^ affect the expression of the neighboring gene *Bacc*, the level of *Bacc* mRNA were measured by RT-qPCR. We found these to be relatively unaffected between control and *Pgk*^ΔE2F^ (Figure S1D).

We concluded that E2F binding sites in *Pgk*^ΔE2F^ are required for the recruitment of E2F/Dp/Rbf to the *Pgk* gene and mediate the full activation of *Pgk* expression throughout development. The effects associated with targeted mutations in the *Pgk*^ΔE2F^ allele are highly specific because the E2F site mutations do not affect the occupancy of E2F/Dp/Rbf at the adjacent site nor significantly alter the expression of the neighboring gene *Bacc*. Given that among the five generated alleles that targeted the E2F sites, the *Pgk*^ΔE2F^ allele had the strongest effect on both recruitment of E2F/Dp/Rbf and gene expression, we selected *Pgk*^ΔE2F^ for further analysis.

### Mutation of E2F sites in *Pgk*^ΔE2F^ lines severely reduced levels of glycolytic and TCA cycle intermediates

PGK catalyzes one of the final steps in glycolysis. The reduction of *Pgk* expression in *Pgk*^ΔE2F^ mutant raises the question whether it leads to an alteration in metabolic homeostasis. To address this question, we measured the steady-state levels of the glycolytic and tricarboxylic acid (TCA) cycle intermediates (Figure 3A). We collected 5 days-old *Pgk*^ΔE2F^ and *w^1118^* control males that were reared in identical non-crowded conditions and determined the levels of selected metabolites by Gas Chromatography – Mass Spectrometry (GC-MS) (Figure 3B). Given that PGK is a glycolytic enzyme we first examined the level of lactate that is commonly used as a readout of glycolytic activity. Lactate pool size was strongly reduced in *Pgk*^ΔE2F^ mutant animals (Figure 3B). The level of dihydroxyacetone phosphate (DHAP) that is interconverted with Glycose-3-Phosphate (G3P) and, thus, can be used as a proxy for the G3P level, was also reduced. Our results suggest that the glycolytic intermediates both downstream of PGK (lactate) and upstream of PGK (DHAP) are reduced. Thus, our findings indicate that the activity of the glycolytic pathway is perturbed in *Pgk*^ΔE2F^ mutant.

**Figure 3:**
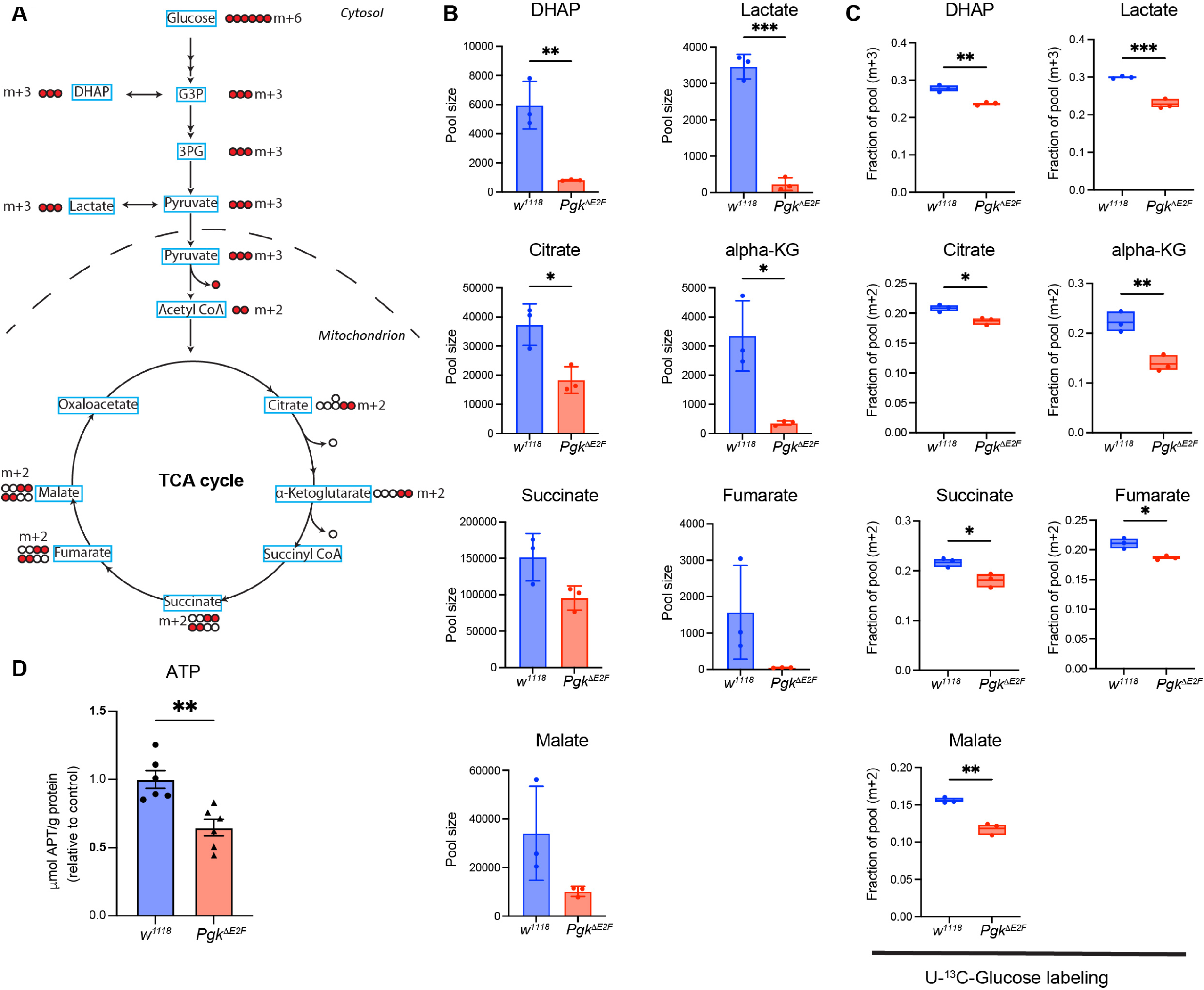
Metabolic changes in *Pgk*^ΔE2F^. (A) Glucose flux in the stable isotope tracing experiment showing the fate of ^13^C from ^13^C- glucose through glycolysis and TCA cycle. (B) Steady-state levels of lactate, dihydroxyacetone phosphate (DHAP), citrate, alpha- ketoglutarate (alpha-KG), succinate, fumarate and malate measured by Gas Chromatography – Mass Spectrometry in *w^1118^* and *Pgk*^ΔE2F^ 5-days-old males. Data presented as bar plot showing Mean ± SD, unpaired t-test: * p < 0.05, *** p< 0.01. (C) Contribution of ^13^C glucose to DHAP, lactate, citrate, α-ketoglutarate, and malate after 12 hours of treatment. M + 3 and M + 2 labeled fractions of pool representing direct flux of ^13^Cglucose into glycolysis and TCA cycle, respectively. Data presented as box plot, whiskers min to max values, line at mean, unpaired t-test: * p < 0.05, ** p< 0.01. n = 3 per group, two independent experiments. (D) ATP levels measured in 5 days old males by luminescence. Total µmol ATP normalized to g protein determined by Bradford Assay. Data presented as bar plot showing mean ± SEM, unpaired t-test, ** p < 0.01, n= 3 samples per genotypes, two independent experiments. Genotypes: *w^1118^* and *w^1118^*; *Pgk*^ΔE2F^ line 12; +

Pyruvate, the end-product of glycolysis, shuttles into mitochondria to fuel the TCA cycle and, thus, serves as a major substrate for the generation of mitochondrial ATP through oxidative phosphorylation. Accordingly, we found a severe reduction in the total levels of several TCA cycle metabolites in *Pgk*^ΔE2F^ mutant, specifically, alpha-ketoglutarate, malate and fumarate and, to a less extent succinate and citrate (Figure 3B). To determine whether low levels of TCA intermediates are due to reduced glycolytic activity in *Pgk*^ΔE2F^ mutant, we used ^13^C-glucose to measure the incorporation of the [U-^13^C] into the glycolytic and TCA cycle metabolites. Adult flies were fed on a ^13^C-glucose diet for 12 hours, then harvested and samples were processed to measure isotope labeled metabolites by GC-MS. Consistent with reduction in steady state levels of glycolytic and TCA cycle intermediates (Figure 3B), the ^13^C labeled DHAP and lactate (m+3 isotopologue) were reduced in *Pgk*^ΔE2F^ mutant compared to control (Figure 3C). Strikingly, there was a corresponding decrease in the ^13^C fraction for several TCA cycle intermediates. As shown in Figure 3C, the ^13^C labeled (m+3) citrate, alpha-ketoglutarate, succinate, fumarate, and malate were significantly lower than in wild type, thus pointing to a reduction in the glucose flux into the TCA cycle. To determine whether abnormal TCA cycle would lead to defects in ATP production, we measured ATP level in extracts prepared from the *Pgk*^ΔE2F^ mutant and matching wild type control males using the bioluminescent assay for quantitative determination of ATP. As shown in Figure 3D, the levels of ATP were strongly reduced in the *Pgk*^ΔE2F^ mutant. The low level of ATP was confirmed in another independent line with the *Pgk*^ΔE2F^ allele (line 23, Figure S2).

Our data show that mutating the E2F sites in the *Pgk*^ΔE2F^ line leads to a reduction in glycolytic activity and disruption of the TCA cycle. Consequently, the generation of ATP is significantly reduced. Thus, we concluded that the loss of E2F regulation on *Pgk* gene expression induces severe metabolic defects.

### The loss of E2F regulation on *Pgk* gene expression leads to mitochondrial defects

The reduction in *Pgk* expression and the changes in the levels of metabolites described above raise the question whether these defects impact the physiology of *Pgk*^ΔE2F^ mutant flies. We reasoned that the tissues expressing high level of *Pgk* mRNA will likely be more sensitive to reduced *Pgk* expression. To determine which organ has the highest expression of *Pgk*, RNA was extracted from different tissues dissected from adult female flies. The level of *Pgk* mRNA across ovaries, dorsal abdomen containing fat body, head and flight muscles was measured by RT-qPCR. Notably, the *Pgk* mRNA was approximately 15-fold higher in flight muscles than in other organs (Figure 4A). Since the expression of the glycolytic genes is coordinately regulated during development, we extended our analysis beyond *Pgk* and compared relative expression of other glycolytic genes among these tissues. Indeed, the levels of *Phosphoglucose isomerase* (*Pgi*), *Phosphofructokinase* (*Pfk*), *Enolase* (*Eno*) and *Pyruvate kinase* (*Pyk*) transcripts were 10 to 20- fold higher in flight muscles than in other tissues, and the increase was almost 100-fold for *Ald1* expression (Figure 4A). These results are consistent with high energy demand in the muscles and the major role of glycolysis in energy metabolism and ATP generation to sustain muscle function (Chatterjee & Perrimon, 2021).

**Figure 4:**
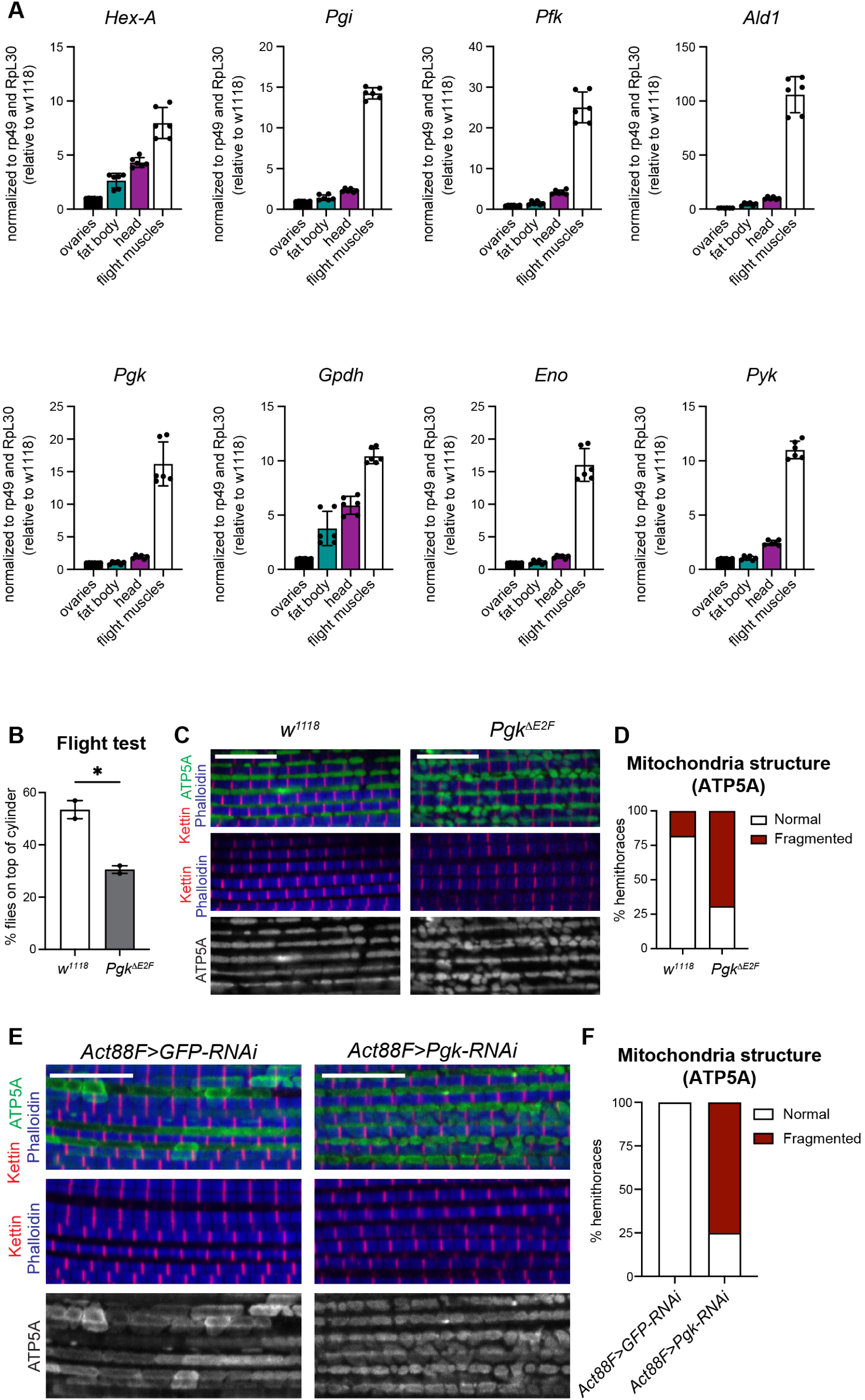
Mitochondrial defects in *Pgk*^ΔE2F^ led to dysfunctional muscles. (A) The expression of the glycolytic genes *Hex-A*, *Pgi*, *Pfk*, *Ald1*, *Pgk*, *Gpdh*, *Eno*, and *Pyk* was measured by RT-qPCR in dissected ovaries, head, flight muscles and dorsal abdomen containing fat body tissue in 2-to 3-days old females. Gene expression was normalized to RpL32 and RpL30 and displayed as fold change relative to control. Scatter dot plots with bars show Mean ± SD, N= 3 independent samples per group (B) Flight ability scored by quantifying the percentage of flies landing on top section of the column. 5- to 7- days-old males were tested. Scatter dot plots with bars depicting mean ± SEM, unpaired t-test, * p < 0.05, N= 196 for *w^1118^* and 158 for *Pgk*^ΔE2F^, two independent experiments. (C, E) Confocal section images of flight muscles in a sagittal view. Hemithorax sections of 5-days old males were stained with Phalloidin (blue), anti-kettin to mark Z-bands on the sarcomere units (red), and anti-ATP5A (green) to label mitochondria structure. (D, F) Defects in mitochondrial structure was visualized by staining with anti-ATP5A as in (C-E) and quantified as the percentage of hemithoraces displaying either normal morphology of mitochondria or fragmented shape (i.e, round). Data are shown as stacked bars. (D) N=11-13 hemithoraces per group and (E) N=11-12 hemithoraces per group Scale 10um Genotypes: (A-D) *w^1118^*, (B-D) *w^1118^*; *Pgk*^ΔE2F^ line 12; +, (E-F) *Act88F-GAL4/Y; UAS-GFP-RNAi;+,* and *Act88F-GAL4/Y; +; UAS-Pgk-RNAi^GL00101^*

The indirect flight muscles are the largest muscles among the thoracic muscles, and their main role is to provide power to fly. We used a flight test assay to quantitively assess muscle function in the *Pgk*^ΔE2F^ mutant flies. In this test, flies are tapped into a cylinder coated with mineral oil. Wild type animals land in the top part of the cylinder, while flies with dysfunctional flight muscle land in the lower part or at the bottom of the cylinder. The flight test revealed that the muscle function is compromised in the *Pgk*^ΔE2F^ mutant flies, as significantly less *Pgk*^ΔE2F^ mutant animals landed in the top part of the cylinder in comparison to control flies (Figure 4B). This result was confirmed with the independent *Pgk*^ΔE2F^ mutant line (line 23, Figure S3A).

To determine whether the weakness in muscle function of the *Pgk*^ΔE2F^ mutants was due to an overall defect in muscle structure, we dissected the indirect flight muscles and stained these with phalloidin and anti-Kettin antibody to visualize sarcomeres, as well as anti-ATP5A antibody to label mitochondria. The wild type muscles displayed a regular array of sarcomeres and the shape of their mitochondria was globular and highly elongated (Figure 4C, left panel). While the sarcomere structure of the *Pgk*^ΔE2F^ mutant muscles was largely indistinguishable from the wild type, the mitochondrial morphology was highly abnormal. Specifically, the shape of mitochondria was highly fragmented and round (Figure 4C-D), which is indicative of mitochondrial disfunction (Avellaneda *et al*, 2021). To ensure that the observed mitochondrial phenotype was specific to the mutations on the E2F binding sites, we confirmed the results using a second independently established *Pgk*^ΔE2F^ mutant line (line 23, Figure S3B-C). Furthermore, defective mitochondria morphology was accurately phenocopied by knocking down the expression of *Pgk* using an *UAS-Pgk*-RNAi^GL00101^ line driven by the flight muscles driver *Act88F-GAL4* (Figure 4E-F). Thus, our data strongly suggest that the reduced expression of *Pgk* in *Pgk*^ΔE2F^ mutant results in dysfunctional mitochondria, consistent with low ATP generation, and thus consequently, lead to defective muscles.

### The *Pgk*^ΔE2F^ allele reduces chromatin accessibility that is accompanied by low expression of metabolic genes

Accumulating evidence suggests that many metabolites serve as co-factors for histone modifying enzymes that play central role in epigenetic regulation (Tsukada *et al*, 2006; Whetstine *et al*, 2006; Xiao *et al*, 2012). One of the striking consequences of the loss of E2F regulation of *Pgk* gene is the severe reduction in several glycolytic and TCA cycle intermediates raising the possibility that these defects may lead to epigenetic changes. To test this idea, we analyzed the global epigenetic landscape in *Pgk*^ΔE2F^ mutant using the Assay for Transposase-Accessible Chromatin with high-throughput sequencing (ATAC-seq). Nuclei from flight muscles of pharate animals were collected and treated with Tn5 transposase followed by PCR to map chromatin accessibility genome-wide (Buenrostro *et al*, 2015). Genomic regions displaying active chromatin regions are open, not densely packed by nucleosomes, and are therefore amplified and sequenced. We obtained high alignment rates with 85% for two replicates samples *w^1118^* and 73% for two *Pgk*^ΔE2F^ samples. Over twenty-five thousand regions peaks were identified as open chromatin regions (Table S1). Each peak was annotated to promoter, 5’UTR, 3’UTR, exon, intron or intergenic region and assigned to the closest TSS using ChIPseeker package (Yu *et al*, 2015) (Table S2).

Next, we compared the chromatin accessibility among sites in common between *Pgk*^ΔE2F^ and control animals using DiffBind (Stark & Brown, 2011). We identified a subset of 513 genomic regions showing a significant reduction in peak intensity in *Pgk*^ΔE2F^ mutants, and only 7 genomic regions with a significant increase (FDR<0.05, Figure 5A-B, S4A). Peaks showing significant changes in chromatin accessibility were annotated to the nearest TSS using ChIPSeeker (Yu *et al*, 2015). Only 3.5% of peaks were in distal intergenic regions while 29.4% were mapped to the gene and 67.1% were located at the promoter site (Figure 5C-D, Table S3). Interestingly, several metabolic genes displayed reduced chromatin accessibility. These included *Pgk*, *Hexokinase A* (*Hex-A*) and *Ald1* from the glycolytic pathway and 1-*Acylglycerol-3- phosphate O-acyltransferase 3* (*Agpat3i)*, *Lipin* (*Lpin*), *minotaur* (*mino*), and *pummelig* (*puml*) from lipid metabolism (Figure 5E-F, Table S3). We noticed that many other metabolic genes, such as *Mitochondrial aconitase 1* (*mAcon1i)*, *Pyk*, *Lipid storage droplet-2* (*Lsd-2*), *Pfk*, *Isocitrate dehydrogenase* (*Idh*), *Acetyl-CoA carboxylase* (*ACC*) and others, had apparent reduction in peak signal albeit with a higher FDR (FDR<0.1, Table S3). Thus, ATAC-seq reveals that chromatin accessibility is broadly reduced in *Pgk*^ΔE2F^ mutant.

**Figure 5:**
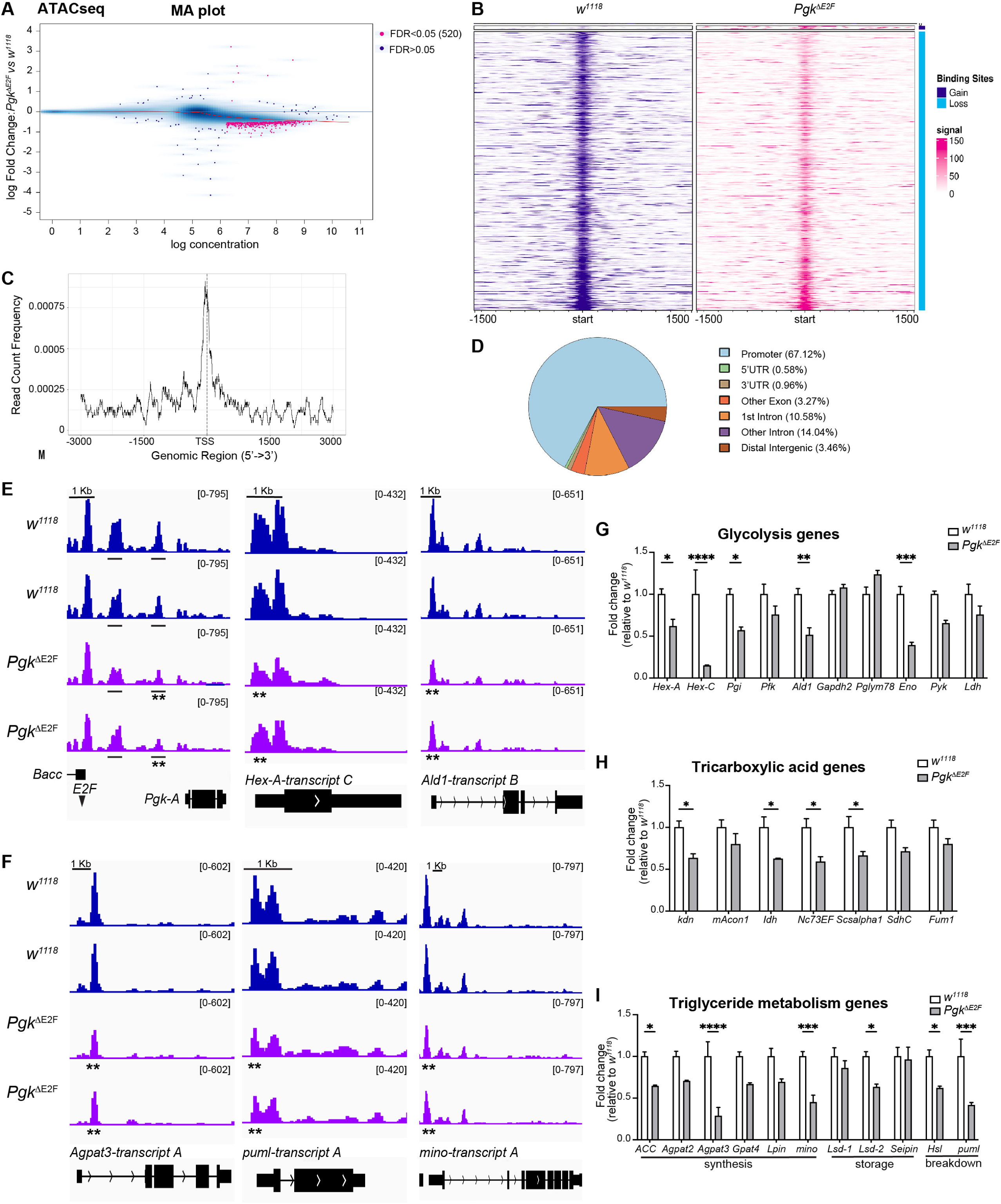
Reduction in chromatin accessibility and gene expression in *Pgk*^ΔE2F^. (A) MA plot representing differential chromatin accessibility analyzed using DiffBind. The common 23,315 sites for ATAC-seq between *Pgk*^ΔE2F^ and control animals were plotted as a blue cloud. An FDR < 0.05 cutoff is symbolized as pink dots. The X-axis values (“log concentration”) represent logarithmically transformed, normalized counts, averaged for all samples, for each site. The Y-axis values represent log 2 (fold change) values in *Pgk*^ΔE2F^ relative to *w^1118^*. Negative log 2 FC values indicate reduced accessibility in *Pgk*^ΔE2F^ (B) Heatmaps showing the enrichment of ATAC reads in a 3000 bp window centered on the summit for each peak. Scale is as indicated in the signal. Only 520 sites with differential chromatin accessibility are included, in which 7 sites show a gain in enrichment in *Pgk*^ΔE2F^ and 513 show a loss (reduction). (C) Distribution of reads for ATAC-seq data in a 6000 bp window centered on the TSS of the gene for each peak. Only the 520 sites with differential chromatin accessibility in *Pgk*^ΔE2F^ are included. (D) Peak annotation pie chart for the 520 sites with differential chromatin accessibility in *Pgk*^ΔE2F^ shows that more than half of the peaks fall into promoter regions. (E-F) Differential chromatin accessibility determined by ATAC-seq visualized with Integrative Genomics Viewer browser for the genomic regions surrounding (E) the glycolytic genes: *Pgk*, *Hexokinase-A* (Hex-A), and *Aldolase1* (*Ald1*), and (F) the lipid metabolism genes: 1-*Acylglycerol- 3-phosphate O-acyltransferase 3* (*Agpat3*), *pummelig* (*puml*), and *minotaur* (*mino*). The most predominantly expressed transcript in adult flies, based on the FlyAtlas 2 profile (Leader *et al*, 2018), is displayed; n = 2 samples/genotype, each track is shown separately for each replicate. Read scales and genomic scales included on top right and top left, respectively. GroupAuto scale was used. ** FDR<0.05 in chromatin accessibility. (G-I) The expression of the metabolic genes measured by RT-qPCR in whole animals staged at pharate (96h APF). (F) glycolytic genes: *Hex-A*, *Hex-C*, *Pgi*, *Pfk, Ald1, Gapdh2, Pglym78, Eno, Pyk, Ldh* (G) TCA genes: *Citrate synthase* or *knockdown* (*kdn)*, mAcon1, *Isocitrate dehydrogenase* (*Idh*), *Aconitase 1* (*mAcon1*), *Oxoglutarate dehydrogenase* (*Nc73EF*), *Succinyl-coenzyme A synthase, alpha subunit 1* (*Scsalpha1*), *Succinate dehydrogenase, subunit C* (*SdhC*) and *Fumarase 1* (*Fum1*) (H) lipid metabolism genes: synthesis (left panel) *Acetyl-CoA carboxylase* (*ACC*), *1-Acylglycerol-3-phosphate O-acyltransferase 2* (*Agpat2*), *1-Acylglycerol-3-phosphate O-acyltransferase 3* (*Agpat3*), *Glycerol-3-phosphate acyltransferase 4* (*Gpat4*), *Lipin* (*Lpin*), *minotaur* (*mino*), storage (middle panel) *Lipid storage droplet-1* (*Lpd-1*), *Lipid storage droplet-2* (*Lpd-2*), *Seipin*, breakdown (right panel) *Hormone-sensitive lipase* (*Hsl*), and *pummelig* (*Puml*). Gene expression was normalized to RpL32 and RpL30 and displayed as fold change relative to control. Bar plots show Mean ± SEM, N= 3 independent samples per group, multiple unpaired t-tests followed by corrected Holm-Šídák method for multiple comparison correction. * p< 0.05 TSS: transcription start site Full genotypes: control *w^1118^* and *w^1118^*;*Pgk*^ΔE2F^ line 12;+

The reduced chromatin accessibility, particularly in promoter regions, indicates potential changes in the expression of the corresponding genes. Therefore, we examined the expression of the metabolic genes that showed reduced ATAC peak intensity in *Pgk*^ΔE2F^ mutant. RNA was isolated from animals staged at pharate and gene expression was measured by RT-qPCR. Remarkably, the levels of metabolic genes mentioned above *Hex-A*, *Ald1*, *Agpat3*, *Lpin*, *mino*, and *puml* were significantly reduced in both *Pgk*^ΔE2F^ mutant lines in comparison to the parental line, *w^1118^* (Figure 5G-I, S4B-C). Interestingly, there was also a reduction in the expression of *mAcon1*, *Pyk*, *Lsd-2*, *Pfk*, *Idh*, *ACC* that had low chromatin accessibility, but with a higher FDR value (FDR<0.1, Table S3). Next, we examined the rest of the glycolytic, TCA cycle and lipid metabolism genes and found that, in total, almost half of them were significantly reduced, while the remaining genes exhibited a similar trend albeit the changes were not statistically significant (Figure 5G-I, S4B-C). Given that the recruitment of Dp and Rbf to the other metabolic targets, such as *Ald1* gene, was unaltered, as shown in Figure 2C-D, we concluded that the changes in gene expression in *Pgk*^ΔE2F^ mutants, described above, are not due to abnormal recruitment of E2F/Dp/Rbf.

To confirm that the changes in the expression of metabolic genes was specific to *Pgk*^ΔE2F^ mutants, we examined the expression of the same panel of genes in heterozygous flies carrying a hypomorphic mutant allele *Pgk^KG06443^*. As expected, there was a two-fold reduction in the *Pgk* mRNA levels in *Pgk^KG06443^*heterozygotes (Figure S4D). However, unlike *Pgk*^ΔE2F^ mutant, in which the *Pgk* expression is downregulated several folds, the expression of glycolytic, and TCA cycle genes was indistinguishable between *Pgk^KG06443^*heterozygous and control animals (Figure S4E- F).

In *Drosophila*, estrogen-receptor related (*ERR*) directs a transcriptional induction that promotes glycolysis, TCA cycle and electron transport chain during pupal development (Beebe *et al*, 2020). Interestingly, *ERR* expression was reduced in both *Pgk*^ΔE2F^ mutant alleles, but not in *Pgk^KG06443^* heterozygotes (Figure S4G-H).

Thus, we concluded that the loss of E2F regulation on *Pgk* gene results in a broad reduction in chromatin accessibility that may reflect changes in global epigenetic profile. Importantly, many metabolic genes associated with decreased accessibility regions were expressed at low levels in *Pgk*^ΔE2F^ mutant.

### The loss of E2F regulation on *Pgk* gene has a broad impact on adult physiology and mimics some aspects of the *Pgk* mutant phenotype

The data described above show that the low expression of *Pgk* in *Pgk*^ΔE2F^ animals severely impacts the levels of glycolytic and TCA cycle intermediates, and the generation of ATP, which impairs the function of high-energy consuming organs, such as muscles. These observations raise the question, what other physiological functions are affected in *Pgk*^ΔE2F^ mutant? Given that previous studies of a temperature sensitive allele *Pgk* allele, *nubian*, have shown that it shortens the life span (Wang *et al*, 2004), we measured the life span of *Pgk*^ΔE2F^ animals and found it to be similarly affected. The median survival of *Pgk*^ΔE2F^ males was significantly reduced in both independent lines carrying the *Pgk*^ΔE2F^ allele compared to *w^1118^* (Figure 6A, S5A). Additionally, we noticed that fewer larvae were hatching from eggs in both independent *Pgk*^ΔE2F^ lines (Figure 6B, S5B), thus, indicating reduced embryonic viability in *Pgk*^ΔE2F^ mutants. Given that this phenotype can be associated with poor oocyte quality (Gandara & Drummond-Barbosa, 2022) we explored oocyte production. We dissected ovaries from both *Pgk*^ΔE2F^ independent mutant animals and found that roughly 30% of their ovaries were smaller than in control animals (Figure 6C, S5C). Next, we stained *Pgk*^ΔE2F^ and control ovarioles with anti-Arm, phalloidin and DAPI to visualize the overall structure of the egg chambers. Remarkably, in agreement with the small size in ovaries, we consistently observed degenerated egg chambers at stage 7-8 in both *Pgk*^ΔE2F^ independent mutant lines (Figure 6D and Figure S5D). Degenerated egg chambers were evidenced by the condensation of the chromatin in nurse cell nuclei at mid-oogenesis, thus indicating that egg chambers were undergoing apoptosis at the onset of vitellogenesis. The choice between egg development or apoptosis is determined by the mid-oogenesis checkpoint that relays on the nutritional status of the female and other hormonal signals (McCall, 2004). When *Pgk*^ΔE2F^ egg chambers passed the mid-oogenesis checkpoint, the uptake of both lipid and glycogen analyzed at stage 10 by staining ovarioles with Bodipy and Periodic acid Schiff, respectively, proceeded normally compared to control egg chambers (Figure S5E-F).

**Figure 6:**
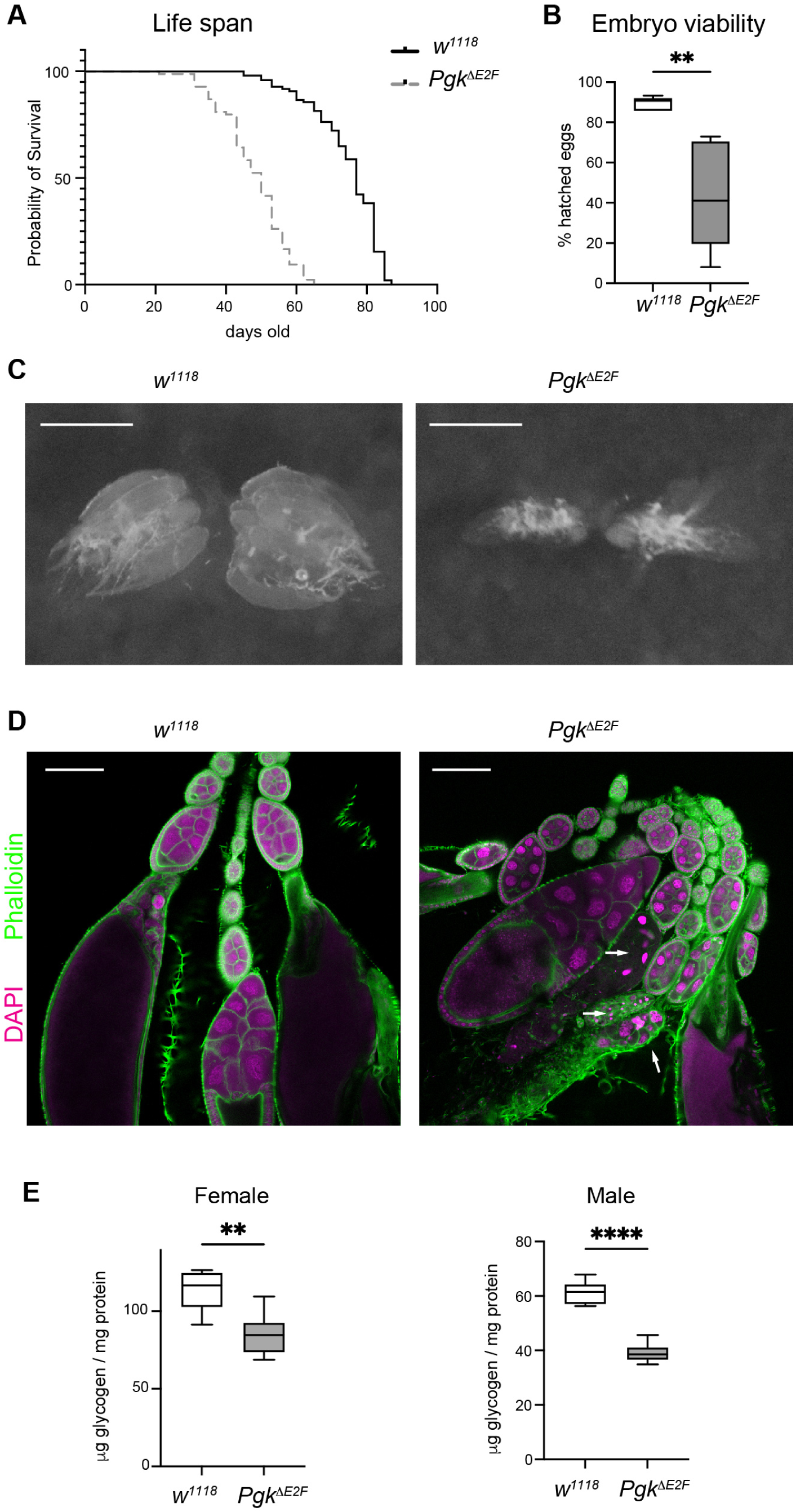
Impact on adult physiology and development in *Pgk*^ΔE2F^. (A) Adult life span determined as survival curves in male flies. Kaplan-Meier analysis, p< 0.0001, median survival = 77 for *w^1118^* and 50 for *Pgk*^ΔE2F^. N=97 and 84 flies per genotype (B) Percentage of hatched eggs quantified as number of first instar larva over number of laid eggs. Data depicted as a box plot, and whiskers represent 5-95 percentile. Mann Whitney test, ** p < 0.01. N=533 eggs for *w^1118^* and 496 for *Pgk*^ΔE2F^ (C) Representative images of the ovaries found in ∼20-40% of females, scale 500 μm. (D) Confocal section images of ovarioles dissected out of 2- to 3-days old females. Ovaries were stained with Phalloidin (green), DAPI (magenta). Condensed and fragmented nurse cell nuclei, indicating degenerated egg chambers (white arrows), found in ∼30% ovaries at mid-oogenesis (around st8), scale 100 μm. (E) 5 days-old females and males were harvested to measure total whole-body glycogen content normalized to protein content. Data are represented as a box plot, and whiskers represent 5-95 percentile, n = 6 per group, two independent experiments were done, unpaired t-test, ** p < 0.01. One representative experiment is shown Full genotypes: control *w^1118^* and *w^1118^*;*Pgk*^ΔE2F^ line 12;+.

The arrest in oogenesis could be indicative of insufficient energetic storage required to meet the biosynthetic demands for egg production (Drummond-Barbosa & Spradling, 2001). As glycogen is a major source of energy storage, we measured its level in the whole animals and normalized the total glycogen content to the protein content. As shown in Figure 6E, the level of glycogen in both males and females were significantly reduced in the *Pgk*^ΔE2F^ mutant compared to the control animals *w^1118^*. This observation was confirmed in another line carrying the *Pgk*^ΔE2F^ mutant allele (line 23, Figure S5G).

Thus, we concluded that the loss of E2F binding sites that contribute to activate the expression of *Pgk* gene broadly impacts animal physiology and leads to shorten life span, reduced embryonic survival, smaller ovaries, degenerated egg chambers and reduced glycogen storage. Collectively, these results illustrate a broad manifestation of the metabolic disbalance in *Pgk*^ΔE2F^ mutant at the organismal level.

## Discussion

E2F regulates thousands of genes and is involved in many cellular functions (Weinmann *et al*, 2002; Xu *et al*, 2007; Korenjak *et al*, 2012; Zappia *et al*, 2019; Cam *et al*, 2004). One of the central unresolved issues is whether E2F function is the net result of regulating all E2F targets or only a few key genes. However, the importance of individual E2F targets cannot simply be deduced by either examining global transcriptional changes in E2F-deficient cells or analyzing ChIP validated lists of E2F targets. The most straightforward approach is to mutate E2F sites in the endogenous E2F targets, yet this was done in very few studies (Tavner *et al*, 2007; Payankaulam *et al*, 2022; Burkhart *et al*, 2010). Genome editing tools provided us with the opportunity to begin addressing this question by systematically introducing point mutations in E2F sites of genes involved in the same biological process. Despite using the same stringent criteria in mining proteomic, transcriptomic and ChIP-seq to select E2F targets, we find that this knowledge is largely insufficient to predict the functionality of E2F binding sites as well as the contribution of E2F in controlling the expression of individual targets. Notably, we showed that among five, similarly high confidence metabolic targets, the loss of E2F regulation is particularly important only for the *Pgk* gene (Table 1). Our findings underscore the value of this approach in dissecting the function of the transcription factor E2F.

**Table 1.**
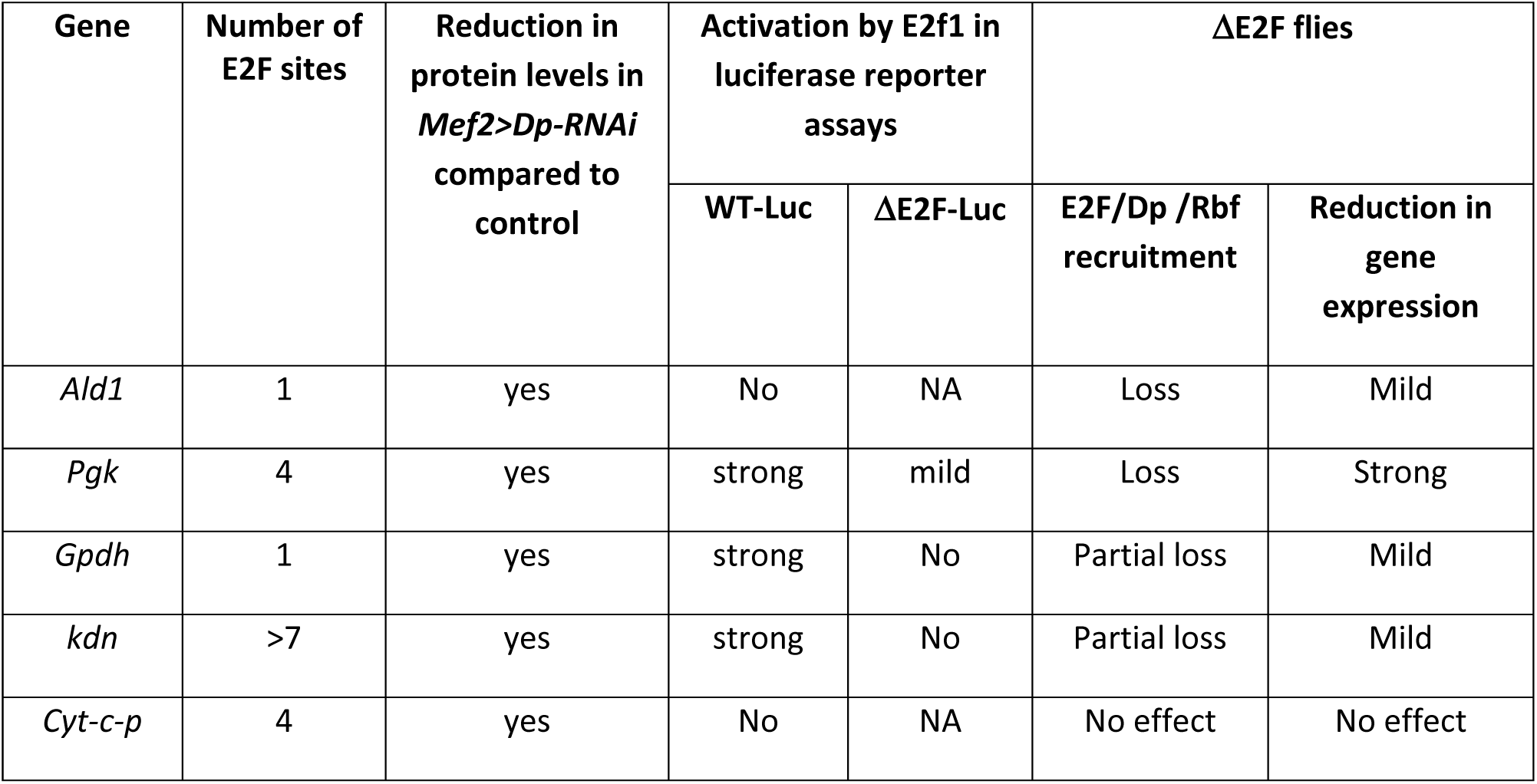
Summary of the effects on mutating E2F sites among five metabolic genes.

The availability of orthogonal datasets for proteomic changes in Dp-deficient muscles combined with genome-wide binding for Dp and Rbf in the same tissue allowed us to select for target genes that should be highly dependent on E2F regulation. Surprisingly, we find significant variation in the recruitment of E2F/Dp/Rbf and the expression of target gene upon mutating E2F site among five metabolic genes (Table 1). Neither the number of E2F sites, their positions nor the response of the luciferase reporter to E2F1 in transient transfection are sufficient to predict E2F regulation of the endogenous targets *in vivo*. Strikingly, although *Cyt-c-p* contained multiple E2F binding sites, the introduction of mutations in these sites had no effect on the recruitment of E2F/Dp/Rbf and the expression of *Cyt-c-p*. For the remaining genes, mutations in the core element of the E2F binding sites reduced the recruitment of E2F/Dp/Rbf and, consequently, the expression of the target gene albeit to a various degree. Our work illustrates that the contribution of E2F in gene expression regulation is highly context- and target-dependent and that one could not simply predict the functionality of E2F binding sites or the importance of a target by relying on existing transcriptomic, proteomic, and ChIP-seq datasets.

In the case of *Gpdh* and *kdn* genes, mutating E2F sites merely results in a partial loss of the binding of Dp and Rbf in the ΔE2F site-edited lines. One explanation for this partial loss in binding is that since the whole animal was used for ChIP-qPCR we cannot distinguish whether the reduction in the recruitment is due to the loss of E2F binding in either a subset of cells or across all cells, thus consistent with a tissue-specific role of E2F. Another possibility is that E2F still retains a weak affinity to the mutant sequence. It is also possible that E2F sites are not the only DNA elements that recruit E2F complexes to chromatin. Indeed, other DNA sequences, such as cell cycle homology region sites, were shown to help in tethering the E2F complex to DNA as a part of the MuvB complex (Müller *et al*, 2014). Additionally, it has been suggested that other DNA binding factors may facilitate E2F binding (Rabinovich *et al*, 2008; Sanidas *et al*, 2022). Overall, our results suggest that, unlike *in vitro* assays with recombinant proteins, mutation of E2F sites does not always fully prevent the recruitment of E2F *in vivo*.

Among five genes that we analyzed here, mutating E2F sites upstream *Pgk* gene completely prevents the recruitment of E2F/Dp/Rbf and results in several fold reduction in the expression of *Pgk* gene. PGK catalyzes conversion 1,3- diphosphoglycerate to 3- phosphoglycerate at the terminal stages of glycolysis to generate one molecule of cytosolic ATP. Decreased *Pgk* levels may lead to a reduction in the production of both glycolytic ATP and pyruvate levels, thus, reducing the major substrate for mitochondrial ATP production. Indeed, neuron specific *Pgk* knockdown markedly decreases the levels of ATP and results in locomotive defects (Shimizu et al., 2020). In another study, flies carrying a temperature sensitive allele of *Pgk*, *nubian*, exhibit reduced lifespan and several fold reduction in the levels of ATP (Wang *et al*, 2004). Thus, altered generation of ATP and shorten lifespan appear to be the hallmarks of diminished PGK function in flies. Notably, the *Pgk*^ΔE2F^ mutant animals show low *Pgk* expression, low ATP levels and die earlier, thus suggesting that in the absence of E2F regulation the normal function of PGK is compromised. This conclusion is supported by our findings that the glycolytic and TCA cycle intermediates are low in *Pgk*^ΔE2F^ mutant.

The loss of E2F function on *Pgk* gene leads to defects in high energy consuming organs, such as flight muscles and ovaries. The morphology of mitochondria is abnormal in *Pgk*^ΔE2F^ and muscles are dysfunctional, as revealed by flight test. The ovaries in *Pgk*^ΔE2F^ females were smaller and contained a high number of degenerated egg chambers at the onset of vitellogenesis. Intriguingly, just like in the *Pgk*^ΔE2F^ mutant animals, occasional degenerating early egg chambers (stage 8 and earlier) were previously described in the *Dp* and *E2f1* mutants (Royzman *et al*, 2002) raising the possibility that this aspect of *Dp* mutant phenotype may be due to loss of E2F regulation on *Pgk* gene. Interestingly, degenerating egg chambers has previously been associated with metabolic defects, including alteration in the TCA cycle intermediates (Rai *et al*, 2022), nutritional status (Drummond-Barbosa & Spradling, 2001; McCall, 2004) and lipid transport (Matsuoka *et al*, 2017). Egg chambers can degenerate before investing energy into egg production, in particular vitellogenesis. Given that multiple signals are integrated during the mid-oogenesis checkpoint, we reasoned that low levels of glycogen in *Pgk*^ΔE2F^ mutant may contribute, at least to some extent, to egg chamber degeneration in *Pgk*^ΔE2F^ mutant. Collectively, our data strongly argue that E2F is important for regulating the function of PGK and the loss of E2F regulation mimics several known phenotypic traits of flies with reduced PGK activity.

Recent studies revealed that metabolic intermediates can function as cofactors for histone modifying enzymes by regulating their activities, and therefore change chromatin dynamics. For example, alpha-ketoglutarate is a co-factor for Jumonji C domain-containing histone demethylases (KDMs), and low alpha-ketoglutarate levels can lead to histone hypermethylation (Tsukada *et al*, 2006; Whetstine *et al*, 2006). In contrast, fumarate and succinate can serve as alpha-ketoglutarate antagonists and inhibit KDMs (Xiao *et al*, 2012). Metabolic analysis of *Pgk*^ΔE2F^ animals revealed a severe reduction in glycolytic and TCA cycle intermediates, including alpha-ketoglutarate, succinate and fumarate, thus, raising the possibility that activities for the histone demethylase are altered in *Pgk*^ΔE2F^ mutants. Another metabolite affected in *Pgk*^ΔE2F^ is lactate that serves as a precursor for a new histone modification known as lactylation that was shown to directly promote gene transcription (Zhang *et al*, 2019). Our findings that the loss of E2F regulation on the *Pgk* gene is accompanied by changes in several metabolites, which can regulate the activity of multiple histone-modifying enzymes, raises the possibility that the epigenetic landscape may change. This idea is supported by altered chromatin accessibility at multiple loci in *Pgk*^ΔE2F^ mutants, as revealed by ATAC-seq. Notably, it is accompanied by a reduction in the expression of several genes, including metabolic genes. The effect was specific to the loss of E2F regulation on the *Pgk* gene because it has been validated in two independent lines carrying the mutation on the E2F sites, while no changes in gene expression was observed in *Pgk* loss of function in heterozygous animals (hypomorph *Pgk^KG06443^*). Interestingly, published microarray analysis for *Pgk^nubian^* animals revealed changes in the expression of genes involved in glucose and lipid metabolism (Wang *et al*, 2004). In sum, our data suggest that abnormal flux through PGK leads to changes in chromatin accessibility and subsequently, transcriptional changes.

Comparison of transcriptomes, proteomes and metabolomes for *Rbf* deficient animals led to unexpected conclusion that very few of these metabolic changes actually corresponded to matching transcriptional changes at direct Rbf targets (Nicolay *et al*, 2013). This result raised the question of the relative importance of transcriptional regulation mediated by E2F/Rb function. Our observation that the loss of E2F regulation of a single glycolytic gene can affect chromatin accessibility and elicit broad transcriptional changes at numerous metabolic genes may provide an explanation for this discordance. Ironically, many glycolytic and mitochondrial genes that are misexpressed in *Pgk*^ΔE2F^ mutants are E2F target genes based on ChIP-seq data. However, the changes in their expression occur without changes in E2F recruitment. Thus, the transcriptional changes observed in the E2F-deficient animals is indeed a complex combination of both direct effects of E2F at the target genes and indirect, yet E2F-dependent effects at other targets as exemplified by *Pgk*. These indirect effects were missed when an entire E2F/Rbf module was inactivated. This finding further highlights the value of mutating E2F binding sites in individual E2F target genes to dissect different tiers of E2F regulation.

## Materials and Methods

### Fly maintenance and stocks

Fly stocks were maintained on standard cornmeal food at 25°C. The WT control strain was *w^1118^* (BDSC 3605) and *Pgk^KG06443^* mutant stock (BDSC 14487), *UAS-Pgk-RNAi^GL00101^* (BDSC 35220), *UAS-GFP-RNAi* (BDSC 41559). The line Act88F-GAL4 (Bryantsev *et al*, 2012) was a kind gift from Richard. M. Cripps. The stocks Vas-Cas9 expressing lines on X chromosome (BDSC 55821) and piggyBac transposase containing CFP marker (BDSC 32070) were used.

Cages were set up to minimize the environmental effect. For all experiments except ATAC-Seq, ChIP-qPCR and RT-qPCR from Figure 2, Figure 4A, and Figure S5D-F), cages were set up with 100 females and 50 males and kept at 25°C. Molasses agar plates with yeast paste were replaced every day. Around 24 h after egg laying, 1st instar larvae were collected, and 40 larvae were transferred to fresh vial and raised at 25°C. The 1 days-old adult flies were transferred into a fresh vial and maintained until 5-day-old flies.

### Identification and modification of E2F binding sites

E2F binding sites were identified using the tool Regulatory Sequence Analysis Tools (RSAT, http://rsat.sb-roscoff.fr/dna-pattern_form.cgi). The E2F degenerated motif WKNSCGCSMM (W:A or T, K:G or T, S:C or G, M: A or C) previously identified through *de novo* discovery motif using ChIP-seq data (Zappia *et al*, 2019) was used. The search region was determined by both Dp and Rbf peaks identified by ChIP-seq data (Zappia *et al*, 2019).

The CGC at the center of E2F binding motif, WKNSCGCSMM, was considered as the core sequence, and was modified from CGC to TAT. In case of multiple tandem E2F sites, the modification was done minimally, but to efficiently disrupt those sites.

### Construction of cell culture luciferase vectors

To generate the luciferase reporter, the E2F-binding site was taken from the regulatory regions of the following genes *Gpdh*, *Pgk*, *Ald*, *Pyk*, *kdn* and *Cyt-c-p*. The regions were PCR amplified from *w^1118^* genomic DNA, and *yw* in the case of *Gpdh*. Primers used are indicated in Table S4. The insert was cloned into the SacI and XhoI sites of the pGL3-HSP70 vector, which is a pGL3 luciferase reporter vector (Promega) with an Hsp70 minimal promoter (Nicolay *et al*, 2011), except for kdn, in which the SacI and NheI sites were used. The pGL3-HSP70-PCNA luciferase reporter was used as a control (Sawado *et al*, 1998).

The mutant fragments for *Pgk*, *Ald*, *Pyk*, *Cyt-c-p* and *kdn* were PCR amplified using the donor plasmids used for *Pgk*^Δ*E2F*^, *Ald*^Δ*E2F*^, *Pyk*^Δ*E2F*^, *Cyt-c-p*^Δ*E2F*^ and *kdn*^Δ*E2F*^ flies, respectively (see details below). The mutant construct for *Gpdh* was made using QuickChange Lighting Site-Directed Mutagenesis kit (Agilent Technologies #210518). The wildtype construct was used a template for mutagenesis, and the manufacturer’s protocol was followed.

The sequence was confirmed by Sanger sequencing. Details regarding the sequence cloned in the luciferase reporters are listed in in the Table S5.

### Cell culture, transfection, and luciferase assay

S2R+ cells were seeded with 60% confluency in a 24-well plate using Schneider’s medium (GIBCO, 2018-03) supplemented with 10% v/v fetal bovine serum (FBS). The next day cells were transfected using X-tremeGENE HP DNA transfection reagent (Roche 06-366-236-001) according to manufacturer’s protocol. About 200 ng of the pIE1-4-E2F1 overexpression construct (Novagen,(Frolov *et al*, 2001)) and 200 ng of the empty pIE1-4 vector (Novagen) were transfected along with the 5 ng of the Luciferase reporters (wildtype and mutant constructs, see details above). About 0.5 ng of the Renilla luciferase plasmid was co-transfected to normalize for transfection efficiency, as a loading control. The reporter assay was performed 48hr post-transfection using dual-luciferase reporter assay system (Promega E1910). The measurements of Firefly and Renilla luciferase activity were made in duplicate in a white, flat bottom, 96 well plate using Synergy H1 microplate reader (BioTek). The volume dispensed was 25 µl with a rate of 250 µL/sec. A 2 second delay was set prior to reading. The integration time was 10 seconds to measure the Luminescence Endpoint with a gain of 135.

### Construction of CRISPR lines

#### I. Injection material preparation

1. Single-guide RNAs (sgRNAs) construct: Targeting sgRNA were cloned by annealed oligos into the pU6-BbsI-chiRNA (Addgene # 45946) plasmid via the BbsI restriction sites. The sgRNA target site in the injecting Drosophila strain, BDSC stock #55821 containing vas- Cas9 on X chromosome, was sequenced and analyzed by Sanger sequencing. Primer sequences are listed in Table S4.
2. Donor plasmid construction. The following components were assembled via Gibson assembly:
  a. Backbone plasmid with negative selection marker. All the pieces necessary for genome editing via Homology Direct Repair (HDR) were inserted via NEBuilder HiFi DNA Assembly Master Mix (NEB #2621) into a plasmid with a negative selection marker that can be used to screen against integration of the entire plasmid into the genome. We used pBS-GMR-eya(shRNA), plasmid #157991 from Addgene. If full plasmid integrates into the Drosophila genome, it results in small eyes and can be used in any line with normal eye morphology. The plasmid was linearized via EcoRV restriction enzyme digestion using EcoRV-HF (New England Biolabs #R3195S) before Gibson assembly.
  b. Modification of interest. The modification of interest (MOI) was located within a range of 18-45bp from sgRNA target site (Chr 2L: 2752555-2752574) for *Pgk*, 161bp from sgRNA target site (Chr 2L: 5943756-5943774) for *Gpdh*, 344bp from sgRNA target site (Chr 3R: 26262086-26262104) for *Ald1*. A total of three modifications of interest (MOI) were located 24bp upstream, 115 bp and 178bp downstream from sgRNA target site (Chr 2L: 16719790-16719808) for *Cyt-c-p*. A total of five modifications of interest (MOI) were located 190bp, 205 bp,264bp,494bp and 573bp from sgRNA target site (Chr X: 6352455-6352473) for *kdn*. MOI was determined by motif searching as indicated in the section *Identification and modification of E2F binding site.* The number of E2F sites and the corresponding edition are shown in Figure 1 and Table S6.
  c. Positive transformation marker. The 3xP3-DsRed marker gene was used to screen for positive integration of the intended modification via HDR. The 3xP3-DsRed cassette from the pScarlessHD-DsRed plasmid (Addgene #64703) is flanked by piggyBac transposition sites (TTAA) that can be used to cleanly excise the entire marker gene after successful integration of the modification via HDR. After excision, the remaining genome sequence was reduced to a single TTAA site. The Scarless Unique primer sequences are Forward: 5’-ATATTGTGACGTACGTTAAAGAT-3’ and Reverse: 5’-GCATTCTTGAAATATTGCTCTCT-3’
  d. Left and right homology arms. The primers used are listed in Table S4. The full sequence of the homology arms is indicated in Table S6.

The Q5 High-fidelity DNA polymerase (New England Biolabs, #M0491) was used for PCR reactions with 30-50 ng of pScarlessHD-DsRed plasmid as template (Addgene #64703) or 50 ng of gBlocks or genomic DNA from the same Drosophila strain (BDSC #55821). Digested product and entire PCR product were run on a 1% agarose gel, and the band of interest was excised from gel and purified using QIAquick Gel Extraction Kit (Qiagen #28706).

All fragments were assembled using the NEBuilder HiFi DNA Assembly Cloning Kit (New England Biolabs, #E5520S) according to manufacturer’s protocol. After incubation at 50°C for 2 hours, it was transformed into NEB 5-alpha competent *E.coli* (NEB #C2987). The sequence of donor plasmid was confirmed by Sanger sequencing using 5 primers to ensure that there are no disabling mutations in the scarless DsRed cassette or regions of the homology arms. Primer sequences are listed in Table S4. Donor plasmid was purified using the Plasmid Plus Midi Kit (Qiagen #12943) prior to injection.

#### II. Injection, screening of transformants, backcrosses and removal of DsRed marker

Injections were performed by Bestgene (www.Thebestgene.com). About 300 embryos of the Vas-Cas9 expressing lines on X chromosome (BDSC 55821) were injected with the sgRNA construct and donor plasmid. Once G0 adults eclosed, flies were individually crossed to healthy virgins or males from a wild type *w^1118^* (BDSC 3605). G1 adults were screened for the expression of 3xP3-DsRed and the absence of eya (shRNA) phenotypes. Positive G1 adults that contained the desired edit were individually crossed to balancer stock. Once lines were established and stable, verification of the anticipated editing/modification was done by PCR analysis and Sanger sequencing. To remove genetic background differences between control, *w^1118^* (BDSC 3605) and genome-edited lines, the backcross with *w^1118^* was performed 6 times, resulting in ∼98.4% synchronism. Precise excision of the 3xP3-DsRed marker cassette from the edited lines was done to achieve scarless genome editing. The piggyBac transposase containing CFP marker on the same allele (BDSC 32070) was used. PCR followed by Sanger sequencing was done to verify that the genome edit was still present after scarless excision.

### RT-qPCR

A total of 3 pharate staged at 96 hours after pupa formation (APF) and 5 third instar larvae were collected per sample. In the case of tissue-specific experiment, females 2-to 3-days old were used and dissections were done in cold- PBS1x. Ovaries and heads from ∼ 10 females were collected per sample. Indirect flight muscles were dissected out of 8 to 10 thoraces per sample. The fat body along with its abdomen were harvested from ∼ 8 females.

Total RNA was purified using 1 mL TRIzol reagent (Invitrogen #15596026). cDNA was made with 1ug RNA using SensiFast cDNA synthesis kit (Bioline, BIO-65053). RT-qPCR was performed using Light Cycler 480 II system (Roche). SensiFAST SYBR No-ROX kit (Bioline, BIO-98005) was used for qPCR reaction, and mRNA levels were normalized to the geometric average of RpL30 and RpL32. All reactions include two technical replicates and three biological replicates. Primer sequences are listed in Table S4.

### Chromatin immunoprecipitation analysis

Twenty 3^rd^ instar larvae were homogenized in buffer A1 (60mM KCl, 15mM NaCl, 4mM MgCl_2_, 15mM HEPES (pH7.6), 0.5% Triton X-100, 0.5mM DTT, 10mM sodium butyrate, and EDTA-free protease inhibitor (Roche, C762Q77)) containing 1.8% formaldehyde, were incubated for 15 min at room temperature, and were quenched with 0.25M glycine for 5 min (Nègre *et al*, 2006). Then the homogenate was washed 4 times with buffer A1, and was washed with lysis buffer (140mM NaCl, 15mM HEPES (pH7.6), 1mM EDTA, 0.5mM EGTA, 1% Triton X-100, 0.5mM DTT, 0.1% sodium deoxycholate, 10mM sodium butyrate, and EDTA-free protease inhibitor (Roche, C762Q77)). Chromatin was resuspended in lysis buffer containing 0.1% SDS and 0.5% N-lauroylsarcosine and was incubated at 4°C for 10min. Then sonication was performed using Branson Sonifier 450 (power 45%, 20 pulses, duration 30sec, interval 59.9sec) on ice. The supernatant was obtained by centrifuging the sonicated chromatin at 4°C for 15min at maximum speed. Immunoprecipitation was carried out overnight with antibodies (mouse anti-Myc(9E10) 1:100, mouse anti-Rbf DX3 and DX5 1:10 each (Du *et al*, 1996), rabbit anti-IgG 1:500, rabbit anti-Dp 212 1:500 (Dimova *et al*, 2003) and Dynabeads protein G (Invitrogen # 10003D). Chromatin precipitate was eluted from Dynabeads with elution buffer 1 (10mM EDTA, 1% SDS, 50mM Tris-Cl pH 8.0) and elution buffer 2 (10mM Tris-HCl pH 8.0, 1mM EDTA, 0.67% SDS) sequentially, and combined eluates were treated with 200ug/ml proteinase K (NEB, P8107S) at 65°C for overnight to reverse crosslinks, and then were incubated with RNAse A (Sigma-Aldrich, 556746) at 37°C for 2 hours. Immunoprecipitated DNA was purified by phenol-chloroform extraction and was precipitated with 100% EtOH containing 0.2M NaCl, and glycogen blue (Invitrogen AM9515).

The qPCR reactions were performed using SensiFAST SYBR No-ROX kit (Bioline, BIO-98005) on the Light Cycler 480 II system (Roche). All reactions include two technical replicates and two biological replicates, except anti-Rbf and anti-myc on kdn gene. Primer sequences are listed in Table S4. Each IP value was first calculated as % input where each IP value was divided by corresponding input value then multiplied by 100. Then it was normalized by Arp53D % input, since Arp53D was considered as a known positive control site where E2F/Rbf binding was previously confirmed. A region near the Mef2 gene was used as a negative control site for E2F/Rbf binding.

### Flight test

The 5-day-old males were used for the flight test as previously described (Zappia & Frolov, 2016b; Zappia *et al*, 2020). Before testing, 25 flies per vial were collected and given 24 hours of recovery time from anesthetization by CO2. The flies were flipped to the 2,000ml cylinder coated with an oil-covered paper. The number of flies attached to each section of the paper were imaged and automatically counted using ImageJ v1.53. A total of 100 flies per genotype were tested. The percentage of flies landing on each section of the cylinder was calculated.

### Immunostaining and confocal imaging

In the case of the flight muscles, thoraces from at least eleven 5-day-old males per genotype were dissected. Thoraces were fixed for 20 min in 4% formaldehyde in relaxing buffer (20 mM Sodium phosphate, 5 mM MgCl2, 0.1% Triton-X, 5mM EGTA, 5 mM ATP), bisected in the sagittal plane, and immediately fixed for an additional 15 min (Weitkunat & Schnorrer, 2014). In the case of ovarioles, ovaries of 5-days-old females were dissected and fixed in 4% formaldehyde in PBS1x for 10 minutes.

Tissues were permeabilized in 0.3% Triton X-100 in PBS, blocked in 2% bovine serum albumin (BSA) 0.3% Triton X-100 in PBS for 1 hour and incubated with primary antibodies overnight at 4°C. Washes for 10 min each were repeated a total of four times in 0.3% Triton X-100 (in PBS) and followed by incubating with Cy3- or Cy5-conjugated anti-mouse and anti-rat secondary antibodies (1:300, Jackson Immunoresearch Laboratories) for 2 hours in 10% normal goat serum and 0.1% Triton X-100 in PBS. After four washes with 0.3% Triton X-100 (in PBS), tissues were stored in glycerol with antifade at 4°C and then mounted. All steps were carried out at room temperature with gentle agitation otherwise indicated.

The primary antibodies were rat anti-Kettin (MAC155, 1:1000, Babraham Institute) and mouse anti-ATP5A (abcam, ab14748, 1:500), mouse α-Armadillo (N2-71A1, 1∶50; Developmental Studies Hybridoma Bank (DSHB). Phalloidin-Atto-488 (1:500, Sigma-Aldrich, 49409) and 4,6- diamidino-2-phenylindole (DAPI, 1:500) were used.

Fluorescent images were acquired using Laser Scanning Microscope 700 (Zeiss LSM700, Observer.Z1) using x20/0.8, x40/1.2 and x100/1.45 objectives. Images were processed using Photoshop (Adobe Systems). Only representative images are shown. All images are confocal single-plane images. Dorsal longitudinal muscles were imaged for flight muscles.

### Glycogen staining

Glycogen staining was performed using the Periodic Acid-Schiff (PAS) kit (Sigma-Aldrich #395B) as follows (Sieber *et al*, 2016). Briefly, ovaries from ten 5-day-old females *w^1118^* and *Pgk*^ΔE2F^ freshly dissected in PBS were fixed in 4% formaldehyde for 30min at room temperature. After washing with water 3 times,10 min per wash, tissue was treated with PAS for 15min. After washing with water, tissue was incubated in Schiff solution for 1 min. After washing with water 3 times, 10 min per wash, the samples were stained with DAPI solution for 5 min and washed again one more time. Ovaries were observed under the Leica Dmi8 microscope and imaged using 10x lens and the color camera DMC2900.

### Stable Isotope labeling

Twenty 5-day-old males *w^1118^* and *Pgk*^ΔE2F-line12^ were used for preparation. After 4 hours of starvation, in which only water soaked in Whatman paper was supplied, flies were fed for 12hrs a mixture of 0.5M 13C-glucose (Cambridge Isotope Laboratories, Inc #CLM-1396-1) and 0.5M unlabeled glucose (Sigma #G5400). Then, flies were flash-frozen in liquid nitrogen until all samples were collected. Flies were grinded using pellet pestle motor with 50% methanol: 50% water followed by freeze-thawing three times. The lysates were centrifuged for 15min at 4°C. The supernatant was transferred to fresh tube, evaporated completely and incubated for 16 hours at room temperature in 30μl of 1:1 Methoxamine (Thermo scientific, TS-45950):Pyridine (Sigma-Aldrich, 270970) followed by derivatization with 70μl of N-tert-Butyldimethylsilyl-N-methyl-trifluoroacetamide (Sigma-Aldrich, 394882) for 1 hour at 70 °C. Derivatized metabolites were injected onto an Agilent 7890 gas chromatograph and networked to an Agilent 5977 mass selective detector. The abundance of the following ions was monitored: m/z 484-487 for DHAP, m/z 261-264 for Lactate, m/z 459-465 for Citrate, m/z 346-351 for alpha-Ketoglutarate, m/z 289-293 for Succinate, m/z 287-291 for Fumarate, and m/z 419-423 for Malate. Data analysis was performed in Data analysis (Agilent) followed by MATLAB GUI for natural abundance correction (Midani *et al*, 2017).

### Life span

Around one hundred 5-day-old males *w^1118^* and *Pgk*^ΔE2F^ were collected. Between 15 and 20 flies per vial were maintained at 25°C. Every 2-4days, flies were transferred to fresh vials without the use of carbon dioxide, and the dead flies were counted at the time of transfer. The percentage of flies that survived over time was analyzed using a Kaplan-Meier plot. At least two independent experiments were carried out per genotype.

### Embryonic lethality

Around 5-day-old *w^1118^* and *Pgk*^ΔE2F^ were collected. Twenty females and ten males were transferred from vials to the cages with on standard cornmeal food and maintained for 2 days at 25°C to make them get used to the environment. Then egg laying plates (molasses) were replaced every hour, 4 times per day for three consecutive days. Scoring was performed 24 hours after egg-laying to allow enough time for larvae to hatch. The egg hatching ratio was determined as the percentage of hatched eggs out of laid eggs. At least two independent experiments were carried out per genotype.

### ATP

The ATP assay was performed following manufacturer’s guidelines and adapted from (Tennessen *et al*, 2014). The six 5-day-old males were homogenized in 100ul of extraction buffer (6M guanidine-HCl, 100mM Tris (pH 7.5), and 4mM EDTA (pH 8)), boiled for 5min, and centrifuged at 14,500 rpm for 5min at 4°C. The supernatant was transferred to a fresh tube. Protein concentrations were measured by the Bradford assay (Bio-Rad 500-0006). ATP levels were measured using the ATP determination kit (Molecular probes #A22066). Samples were diluted 1/200 in reaction buffer prior to measurement. The luminescence was measured in duplicate using a Synergy H1 microplate reader (Biotek). Ten μL of each sample, std and blank (H20) was loaded in a white opaque 96-well plate (Corning) followed by 100 μL of ATP std reaction buffer to each well using a multichannel pipette. Luminescence was immediately measured at 28C using an integration time of 5 sec. Each experiment was run with 5 independent samples. The ATP level was normalized to total protein concentration.

### ATAC-Seq

Sample preparation: At least 40 pharates per sample staged at 96 h APF were dissected in chilled PBS1x 0.1%BSA. The flight muscles were collected from the thoraces. Two independent replicates per genotype were processed. Protocol as in (Buenrostro *et al*, 2015) was followed with some modifications (Merrill *et al*, 2022). Briefly, the nuclei were extracted by grinding tissue using a pellet pestle motor in chilled lysis buffer (10 mM Tris-HCl pH 7.5, 10 mM NaCl, 3 mM MgCl2, 0.1% Tween-20, 0.1% Nonidet P40, 0.05% digitonin (Sigma 300410-250MG), and 1% bovine serum albumin) and further minced using a glass douncer with a type-B pestle and adding TritonX-100 to reach final concentration 0.5%. Nuclei suspension was centrifuged at 1000 g for 10 min at 4C and resuspended in 225 ul wash buffer (10 mM Tris-HCl pH 7.5, 10 mM NaCl, 3 mM MgCl2, 0.1% Tween-20 and 1% bovine serum albumin). A volume of 300 μl of Percoll solution 9:1 (900 μl Percoll (cytiva 17089101) + 100 μl PBS 10x) was added to nuclei, mixed and centrifuged 20,000 x g for 30 minutes at 4°C to remove myofibrils from the nuclei suspension (Hahn & Covault, 1990). Tagmentation Master Mix was added to the pellet and reaction was incubated in a themomixer at 600-800 rpm set at 37C for 30 min as indicated by manufacturer (ATAC-Seq kit, 53150, Active Motif).

Making libraries and sequencing: Libraries were made as in ATAC-Seq kit (Active Motif, 53150) with some modifications. Briefly, tagmented DNA was purified, PCR amplified with i7 and i5 indexed primers (8 cycles). Double sided size selection using SPRI clean-up (0.6X) was done to select for 100-400 bp range. A second PCR reaction (8 cycles) was followed by PCR clean-up with AMPure XP (Agencourt A63880) bead solution (1.5X the sample volume). The pattern of DNA fragment size was analyzed with a 2% low melting agarose gel (NuSieve GTG agarose, BMA 50081). Gel band within 100-250bp was cut and purified using Zymoclean Gel DNA recovery kit/ DNA Clean & Concentrator-5 (Zymo Research). If needed, a third PCR was set up (5 cycles) to increase the yield. Quality of libraries was assessed using Agilent TapeStation with High Sensitivity D5000 ScreenTape. Libraries were pooled and submitted for sequencing on NovaSeq 6000 SP flowcell with paired-reads 2x150 at the DNA services at University of Illinois Urbana Champaign.

Analysis of ATAC-seq data: Fastq files were generated and demultiplexed with the bcl2fastq v2.20 Conversion Software (Illumina). Reads in fastq files were trimmed to remove adapters using cutadapt 2.3 with Python 3.6.10. Paired reads were aligned using Bowtie2 (v2.4.1) to *Drosophila melanogaster* genome version BDGP6.32 (Ensembl version 105) allowing up to 1 mismatch (Langmead & Salzberg, 2012). The BAM files were generated from SAM files using samtools view (v.1.9)(Li *et al*, 2009). Aligned BAM files were sorted, and PCR duplicates were marked using Picard tools MarkDuplicates program (https://broadinstitute.github.io/picard/). Peak calling was done using MACS2 v2.2.7.1 with parameters BAMPE -g dm --keep-dup 1 -q 0.01 (Zhang *et al*, 2008). Differential accessible peaks were determined using DiffBind_3.2.7 (Stark & Brown, 2011)and R version 4.1.0. A total of 23,315 sites in common among all 4 samples were identified using parameter summit = 50. Data were normalized using background method (csaw package,(Lun & Smyth, 2015)). Briefly, genome is divided into large bins (10,000 bp) of reads across the genome, and it contains mostly “background” reads. Peaks were annotated using annotatePeak tool in ChIPseeker v1.5.1 (Yu *et al*, 2015) and and R version 4.1.0, using the TxDb.Dmelanogaster.UCSC.dm6.ensGene. The annotation assigns whether peak is in promoter, 5’UTR, 3’UTR, exon, intron or intergenic region. The TSS (transcription start site) region, was defined from -1kb to +1kb. The sorted and indexed bam files were converted into bigwig files using the bamCoverage tool in the Deeptoolssuite with parameter - normalizeUsing=RPKM. The normalized bigwig files were uploaded to Integrative Genomics Viewer IGV_2.12.3 to visualize peaks across all genotypes and replicates. FlyBase release FB2022_05 was used to browse gene function and summary (Gramates *et al*, 2022).

### Glycogen content

Seven 5-days-old females and males were harvested in separate tube to measure glycogen (Tennessen *et al*, 2014). Briefly, animals were homogenized in 100 μl chilled PBS 1x using pellet pestle motor. An aliquot was stored at -80°C to measure protein content. Then, all samples were heat-treated at 70°C for 10 min and centrifuged for 3 min at 4°C at maximum speed. Supernatant was transferred to a fresh tube and stored at -80°C. Once all samples were collected, these were diluted 1:8 in PBS 1x for the assay and an aliquot was transferred to two wells. One well was incubated with the amyloglucosidase (Sigma A1602) and the other well with PBS 1x. The plate was kept at 37°C for 1 hour. Total amount of glucose was determined by adding 100 μl of glucose (HK) assay reagent (Sigma, G3293). Plate was incubated for 15 min. Absorbance at 340mm was read in the Plate reader BioTek Epoch. Both glycogen and glucose standard solutions were used as standards to determine the glycogen concentrations for each sample. Values of free glucose in the untreated samples was subtracted from the treated one. The total content of glycogen was normalized to content of soluble protein measured using the Bradford assay (Bio-Rad 500–0006) with BSA standard curves.

Samples were measured twice, and a total of six independent samples were measured per genotype. A total of three independent experiments were done.

### Statistical analysis

Statistical analysis was done using GraphPad Prism v9.4. Details regarding data presentation and statistical analysis were included in legends to Figures.

### Data and software availability

ATAC-seq data from this publication has been deposited to the GEO database (https://www.ncbi.nlm.nih.gov/ geo/) and assigned the identifier Series GSE217225 (https://www.ncbi.nlm.nih.gov/geo/query/acc.cgi?acc= GSE217225)

## Acknowledgment

We thank Nick Dyson for critical reading of the manuscript and Kostas Chronis for advice with ATAC-seq experiments and analysis. We thank Adams Didier for making the *Gpdh-WT-luc* construct for luciferase reporter. We are grateful to R.M. Cripps for sharing fly stock *Act88F- GAL4*. Other stocks were obtained from the Bloomington Drosophila Stock Center (NIH P40OD018537). The α-Armadillo (N2-71A1) antibody was obtained from the Developmental Studies Hybridoma Bank, created by the NICHD of the NIH and maintained at The University of Iowa. We are grateful to Flybase for online resources on the Database of Drosophila Genes and Genomes. This work was supported by NIH grant R35GM131707 (to M.V.F.)

## Author contribution

Conceptualization, M.P.Z. and M.V.F.; Methodology, M.P.Z, and Y-J.K.; Investigation, M.P.Z., Y-J.K., A.W., I.L., and H.M.L; Software, A.B.M.M.K.I.; Visualization, M.P.Z.; Supervision, J.K., and M.V.F.; Writing – Original Draft, M.P.Z. and M.V.F.; Writing – Review & Editing, M.P.Z., A.W., H.M.L., J.K. and M.V.F; Funding Acquisition, M.V.F.

## Supplemental Tables

**Table S1: Peak Calling**

**Table S2: Annotation**

**Table S3: DiffBind_Annotated**

**Table S4: Oligonucleotides**

**Table S5: Sequences of inserts used in luciferase constructs**

**Table S6: Sequences of homology arm gBlocks used to assemble donor plasmid for construction of CRISPR lines**

